# Mutational destabilisation accelerates the evolution of novel sensory and network functions

**DOI:** 10.1101/2023.11.23.564566

**Authors:** Yuki Kimura, Shigeko Kawai-Noma, Daisuke Umeno

## Abstract

Binding-induced folding^1–4^ (BIF) is a promising mechanism that can be used to rapidly convert binders into sensors/regulators without allosteric design. Here we showed that allosteric regulatory proteins AraC can acquire BIF mechanism without compromising their inherent allosteric mechanisms, with high frequency upon mutations. This opened an opportunity to compare the evolutionary capacity of the allosteric and non-allosteric modes of a specific sensory protein. We found that AraC evolved novel sensory function far more rapidly in BIF mode than in allosteric mode. This newly acquired (non-allosteric) sensory function is distinguishable both in its response logic and in sensitivity from original (allosteric) one, and they can be operated simultaneously, independently, and cooperatively, allowing the construction of complex regulatory networks behaviours such as a selective NIMPLY/OR converter and width-tuneable band-pass filter. Together with its high frequency of emergence, BIF can be an overlooked evolutionary driver of the invention of novel biosensors and complex regulatory networks in nature and laboratory.

## Main

Metabolic networks achieve high efficiency, robustness, and dynamic responsiveness to environmental changes through the placement of multiple allosteric regulators in key positions ^5–8^. Each of these regulators has evolved high-fidelity response to the target molecules though a sophisticated allosteric mechanism: to selectively recognise the target molecules in the presence of numerous metabolites, allosteric regulators require a structural transition associated with binding, thereby preventing false-responses to the non-specific interaction with off-target molecules^9,10^.

On the contrary, an over-reliance on allosteric regulators could lead to a reduction in the evolvability of metabolic networks. New allosteric regulators can only be created when a specific binding interface and a mechanism to transmit the new interaction into a clear structural transition have emerged simultaneously^11,12^. This is in addition to the constraint that metabolic regulators fulfil an original function within the existing network. Despite these constraints, new functions are constantly being created in nature, both at the component level and at the network level^13,14^. We are interested in how natural metabolic systems attain functional robustness and evolutionary plasticity, apparently contradictory traits.

We suggest that binding induced folding (BIF) is a possible mechanism for searching for novel sensory and network functions. BIF has originally received attention as a novel and convenient strategy for converting binder into sensors^1–4^. Instead of conducting transition from an inactive structure to an active structure triggered by binding with the target molecule, BIF-sensors exert their response by folding facilitated by interaction with the target molecules.

In this study, we questioned our hypothesis that BIF could be the key features that accelerate exploration of new sensory functions and complex network functions. Upon random mutagenesis, allosteric regulator AraC was converted into binding-induced folder with surprising frequency, without compromising its original function as allosteric regulators. We demonstrate that sensor functions in this BIF mode can access novel sensory functions far more frequently than those operating in the allosteric mode. The novel sensor functions obtained in the BIF mode can work independently and simultaneously with the functions of the allosteric mode in a single cell, and their cooperation has yielded complex network functions in a predictable way and with a minimal set of components. Collectively, mutation-induced destabilisation^15–21^, which has been regarded as a mere evolutionary constraint, can be a largely overlooked accelerator of protein evolution, through which mere molecular binding turns selectable both to the evolving metabolic network and to synthetic biologists.

### AraC-inducing systems

AraC, an L-arabinose-responsive transcription factor^22^, is a classic example of allosteric switches. Without L-arabinose, it forms a dimer and binds to the I1 and O2 sites on the target DNA to form a DNA loop that prevents RNA polymerase from accessing the promoter. Upon binding with L-arabinose, AraC undergoes a conformational change, triggering the re-positioning of the O2-bound AraC to the I2 site, which allows the recruitment of RNA polymerase to activate transcription initiation from the P_BAD_ promoter. Hence, L-arabinose is the allosteric effecter that triggers this dynamic negative-to-positive regulation, enabling high signal-to-noise transcription control (**Fig. 1a**).

**Fig. 1.**
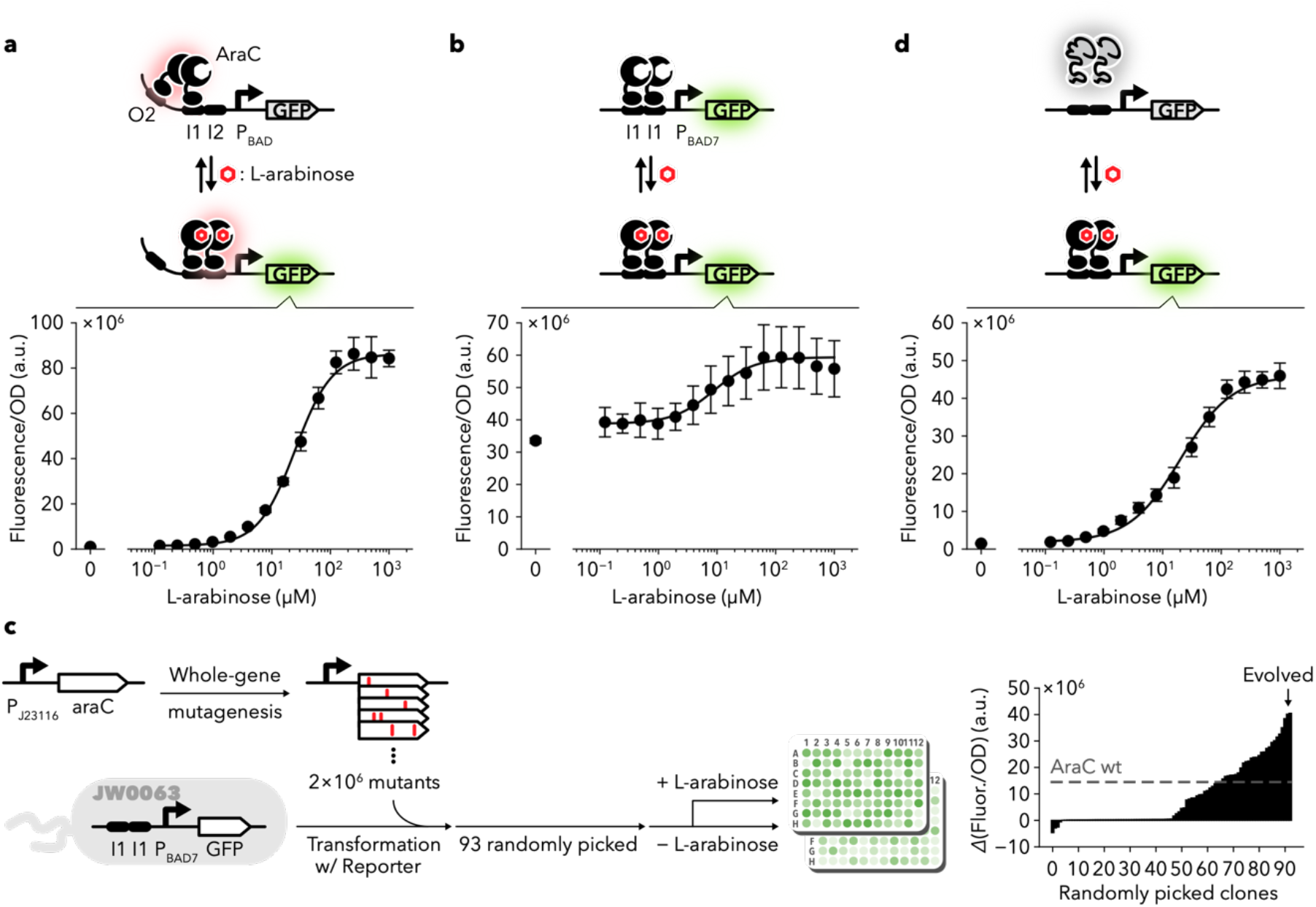
Arabinose-induction systems via allosteric and BIF modes. **a**, Natural AraC/P_BAD_ system (allosteric mode). In the absence of L-arabinose, AraC binds to the I1 and O2 sites to repress transcription. Binding to L-arabinose causes conformational changes in the AraC dimer, which then binds to I1 and I2 sites and activates transcription from the P_BAD_ promoter via interactions with RNA polymerase, inducing the reporter gene (GFP) expression. **b**, AraC/P_BAD7_ system. P_BAD7_ promoter contains tandemly repeated AraC binding site I1, which exhibits the strongest binding affinity for AraC. AraC always binds to the I1-I1 region and activates the transcription of the reporter gene, irrespective of the presence of its ligand L-arabinose. **c**, Workflow for the functional tuning of the AraC/P_BAD7_ system. The grey dotted line represents the function of wild-type AraC. The comparing data in the allosteric and BIF modes is shown in **Supplementary** Fig. 1. **d**, AraC/P_BAD7_ system using AraC mutant (BIF mode). In the absence of L-arabinose, the AraC mutant cannot fold by itself and fails to activate transcription. Mutations found in this mutant, Rndm-B2, are provided in **Supplementary Table 1**. The data shown in **a**,**b**,**d** corresponds to the mean ± s.d. of three biological replicates.

Reeder and Schleif showed that binding of L-arabinose is not a prerequisite for AraC-induced transcription activation when the O2 site is removed^23^. They tested a series of P_BAD_ promoter variants and found an interesting variant termed P_BAD7_. This variant lacks the O2 site and has two tandemly repeated I1 sites, one of which overlaps the −35 box (**Extended Data Table 1**). Because AraC dimer has the highest binding affinity for I1^24^, it occupies the I1-I1 region of P_BAD7_ and strongly activates transcription initiation irrespective of the presence of L-arabinose (**Fig. 1b**). We assumed that this AraC/P_BAD7_ system, which behaves as super-activator (always-ON), can be used as a decent switcher via random mutations in the *araC* gene. This approach is based on our previous observation^4^, where a surprisingly high fraction (∼20%) of the LuxR variant pool, created by random mutagenesis of a super-activator LuxR mutant^25^, exhibited significantly elevated stringency by moderate destabilisation.

We constructed an AraC library using error-prone PCR^26^. Then, we randomly picked 93 variants and examined their ability to activate the P_BAD7_ promoter using a fluorescent reporter in the presence/absence of L-arabinose. While approximately 50% of the variants behaved as super-activators, almost 20% of them showed a significantly improved dynamic range in their response (**Fig. 1c)**. Among the 93 randomly picked variants, we selected five AraC variants that could induce the P_BAD7_ promoter with as high a stringency as the natural AraC/P_BAD_ system (**Fig. 1d**).

Due to their high frequency of emergence, these AraC variants with switching properties might be destabilised by mutations so that they become dependent for stabilisation on the interaction with L-arabinose, as was the case for LuxR^4^. This high on/off ratio of the AraC/P_BAD7_ system is not the result of optimizing regulator expression^27^. Although randomisation of ribosome binding site significantly expands the expression level of wild-type AraC and the maximum output value varied greatly, the on/off ratio of these variants barely changed (**Extended Data Fig. 1**).

We found that certain AraC variants stringently induced both P_BAD7_ and P_BAD_, indicating a moderate decrease in their folding stability without losing the allosteric regulation. A few AraC mutants behaved as good switchers only for P_BAD7_ (**Extended Data Fig. 2**), indicating that the mutations might have caused a structural destabilisation, disrupting the allosteric mechanism.

### Evolving agonistic response to antagonist

While agonists trigger the response of sensor/receptor proteins, antagonists inhibit this response. Although their effects are opposite, the binding of antagonists and agonists to the receptors for their effects is similar. Since BIF can be used to detect stabilisation by a binding event, BIF-based biosensors might respond to an antagonist in the same way as to an agonist.

The molecular structure of D-fucose, a known antagonist for AraC^28^, resembles that of the cognate ligand of AraC, L-arabinose, with a methyl group attached to the carbon at position 5 (**Extended Data Fig. 3**). Although D-fucose binds to AraC, it does not trigger a conformational change of AraC, thereby locking its configuration in the ‘off’ mode (bound to O2 and I1). Indeed, no D-fucose response was observed in the AraC/P_BAD_ system, whereas a slight increase in transcription was observed in the AraC/P_BAD7_ system upon D-fucose binding (**Extended Data Fig. 3)**.

We investigated the frequency of the AraC variants that are induced by D-fucose from the AraC library created by random mutagenesis as shown in **Fig.1c** (**Fig. 2a**). Here again, approximately 20% of the 93 randomly picked AraC mutants exhibited a D-fucose response (**Fig. 2b**, lower), which is consistent with the frequency of switching mutants to L-arabinose (**Fig. 1**). In contrast, only one mutant was found to induce P_BAD_ (**Fig. 2b**, upper).

**Fig. 2.**
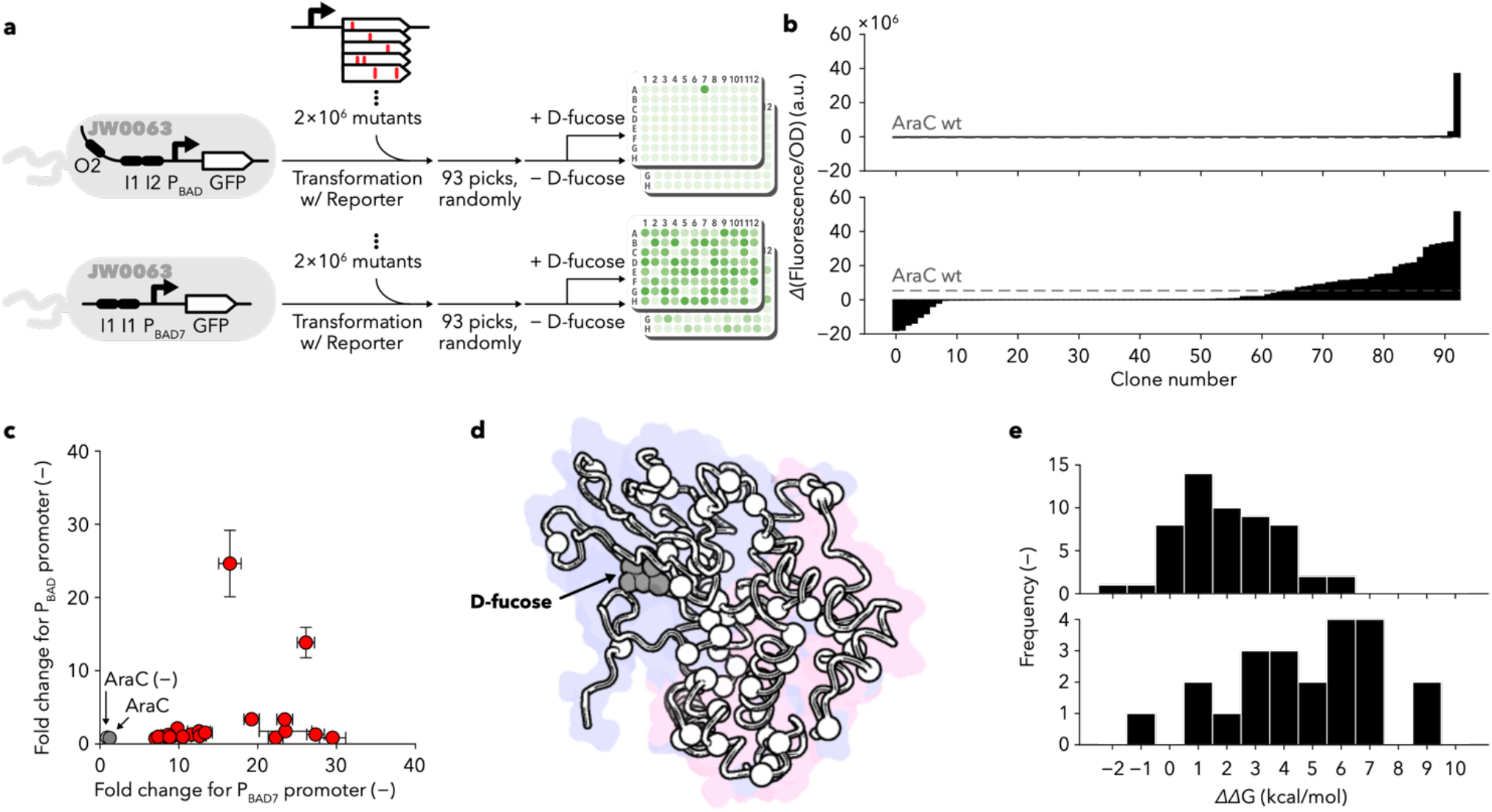
Emergence of antagonist-responsive AraC mutants. **a**, The experimental workflow for searching for D-fucose responders in allosteric and BIF modes. **b**, Fitness landscapes of AraC for the allosteric (upper) and BIF (lower) modes, respectively. The functions of the 93 AraC mutants and the wild type are represented by the solid black and dotted grey lines, respectively. These experiments were performed as a single measurement. **c**, Comparison of the switching modes of the 22 mutants showing ≥ 5-fold response to D-fucose in AraC/P_BAD7_ (BIF mode) (as shown in **c** and Supplementary Fig. 2). Each value corresponds to the mean ± s.d. of three biological replicates. Red, AraC mutants; grey, wild-type AraC; white, AraC (−). **d**, Structural mapping of the mutations found in the 22 D-fucose responsive mutants based on superposition of the structure predicted by AlphaFold2 (ID: AF-P0A9E0-F1-model_v2) and the crystal structure of the ligand binding domain of AraC bound to D-fucose (PDB ID: 2aac). The white spheres indicate the alpha-carbons of the amino acids where the mutations were introduced. The blue and red backgrounds represent the sugar-binding and DNA-binding domains, respectively. **e**, Predicted stability effects of 55 individual mutations (upper panel) found in the 22 mutants (lower panel). The Gibbs energy change (ΔΔG) was calculated by FoldX. The data are presented in histograms with 1 kcal/mol intervals, ranging from −2 to 10 kcal/mol.

We recovered 22 mutants that exhibited a five-fold or greater D-fucose response in the AraC/P_BAD7_ context and examined their response to the P_BAD_ promoter. Most of them failed to induce transcription at P_BAD_ (**Fig. 2c**). Sequence analysis of the D-fucose-dependent P_BAD_ activator AraC variant AraC carried I46V mutation, which might be responsible for the allosteric response to D-fucose. P_BAD7_ behaved as a super-activator, i.e. wild-type-like (**Extended Data Fig. 4**), indicating this mutant acquired novel conformational changes for D-fucose without compromising stability.

Genotyping D-fucose-responsive mutants revealed amino acid substitutions with little bias and without any mutational hot spots (**Fig. 2d** and **Extended Data Fig. 5**). We used the FoldX algorithm^29^ to predict stability changes (*ΔΔ*G) due to mutations. We found that the stability change for all 55 unique mutations was distributed with a peak value of 1 kcal/mol and many mutations tended to decrease the stability (**Fig. 2e**, upper). Furthermore, the data for each mutant showed that most of the mutants were highly destabilised **(Fig. 2e**, lower). Since the binding affinity of AraC for D-fucose is similar to that for L-arabinose (apparent K_d_ of 6 × 10^−3^ M)^30^, it might have acquired its agonist response by moderately reducing its structural stability through random amino acid mutations to become a binding-induced folder. In summary, binding-induced stabilisation is unique because it does not distinguish between agonists and antagonists. Moreover, it allows the development of rapid antagonist-responsive sensors with significantly higher frequency (in this case, 20-fold) without the need to reconstruct the allosteric mechanism.

### Searching for novel sensor functions

Because redesigning allosteric mechanisms is not required, we hypothesised that there is a higher probability that sensors based on binding-induced stabilisation can perform newer and more diverse functions than allosteric sensors.

Unlike D-fucose (antagonist), D-galactose does not bind to AraC^31^. Structural analysis suggested that the sugar-binding pocket formed by F15, M42 and I46 on AraC fits the methyl group of D-fucose (**Fig. 3a**), but fails to accommodate the bulky C5-hydroxymethyl group of the D-galactose^32^.

**Fig. 3.**
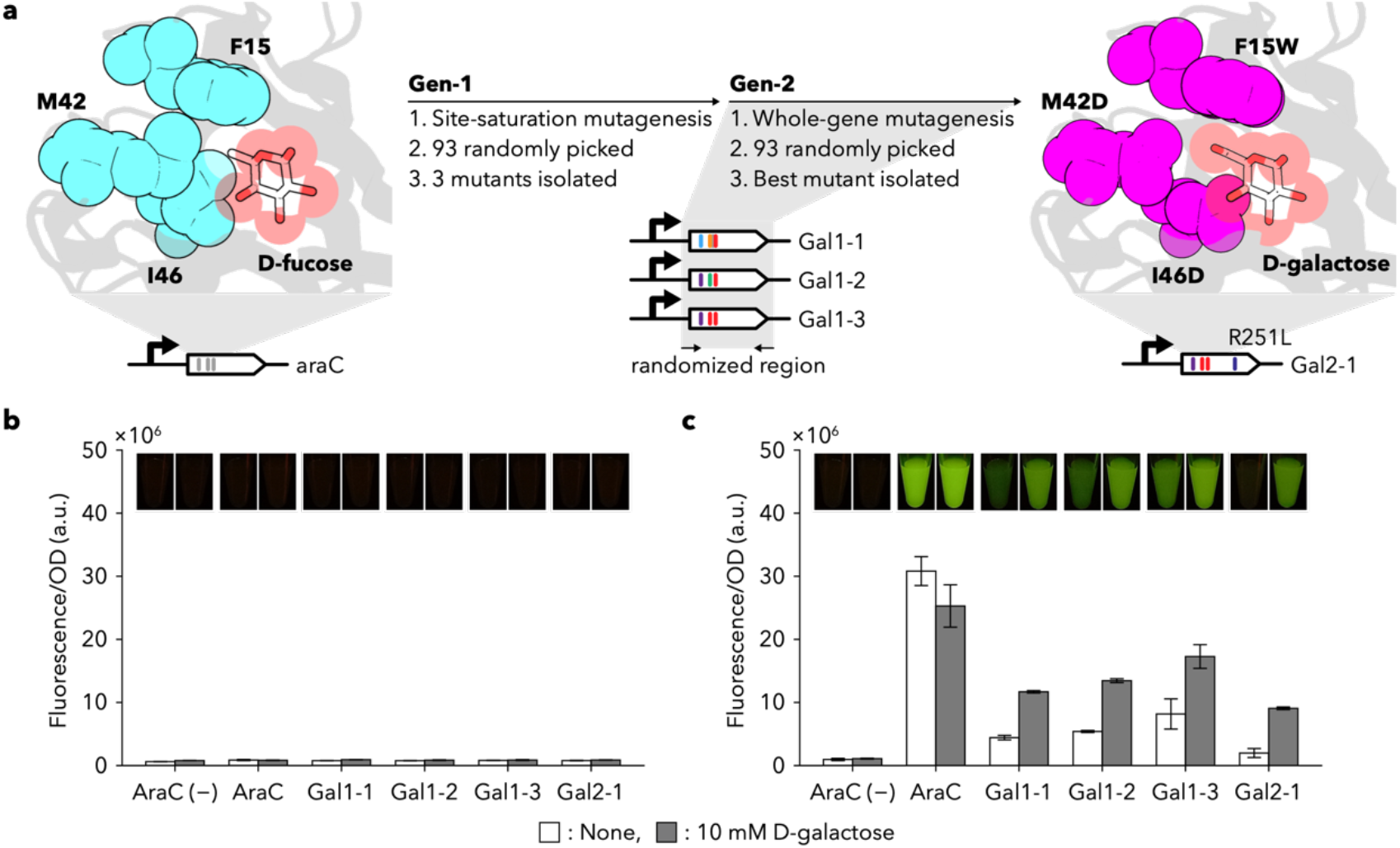
Directed evolution of AraC to D-galactose sensor. **a**, The three residues in contact with the methyl group of D-fucose (shown as a stick) were simultaneously randomised and selected for fluorescence in the presence of D-galactose. Three positive mutants were pooled and subjected to whole-gene random mutagenesis for increased stringency, leading to the isolation of quadruple mutants AraC_F15W, M42D, I46D, R251L_. **b**,**c**, Response of AraC and its variants to D-galactose in allosteric (**b**) and BIF modes (**c**), respectively. Each value corresponds to the mean ± s.d. of three biological replicates. After dispensing 200 µL of cell culture into 1.5 mL microtubes, fluorescent images were captured using Gel Ninja. Sequences found in D-galactose-responsive mutants are provided in **Supplementary Table 2**.

To obtain AraC mutants that bind and respond to D-galactose, we simultaneously randomised these three amino acids, F15, M42 and I46. From the 93 randomly picked AraC variants, we found three D-galactose-responsive mutants (**Extended Data Fig. 6**a), which were pooled and subjected to random mutagenesis by error-prone PCR. Again, among the 93 mutants randomly selected from the mutant pool, one exhibited a D-galactose response with a decent stringency (**Extended Data Fig. 6**b). Note that, this second-generation mutant (Gal2-1) is just the best among the 93 randomly selected variants.

All four mutants with measurable D-galactose responses exhibited an always-off behaviour toward the P_BAD_ promoter (**Fig. 3b**). Thus, any combinatorial mutations that conferred D-galactose response in the BIF mode (**Fig. 3c**) did not invoke the allostery. Also, it indicates that the evolution of AraC as a D-galactose responder requires additional mutations that establish the intra-molecular interactions needed for allosteric switching.

AraC is one of the most extensively engineered sensor proteins. Seminal work by Cirino and colleagues created AraC variants that can allosterically respond to D-arabinose^33^, mevalonic acid^34^, triacetic acid, lactone^35,36^, ectoin^37^, vanillin^38^, salicylic acid^38^ and orsellinic acid^39^ by engineering the binding specificity of AraC. This remarkable set of functions was achieved via computational design and ultra-high throughput FACS sorting. The frequency of the emergence of these mutants approached 10^−6^.

Based on the available literature, we selected a salicylic acid-responsive AraC mutant, named Sal4 (ref.^38^), which was obtained through FACS selection at a frequency of 10^−6^ from a large library after simultaneously randomising five ligand-binding residues. Sal4 has five amino acid substitutions P8V, T24I, H80G, Y82L and H93R (**Figs. 4a and c**). We created 32 AraC variants with all combinations of these five mutations (n = 2^5^) and examined the changes in fluorescence upon the addition of salicylic acid, both in the allosteric and the BIF modes (**Figs. 4b, d and e, Extended Data Fig. 7**a).

**Fig. 4.**
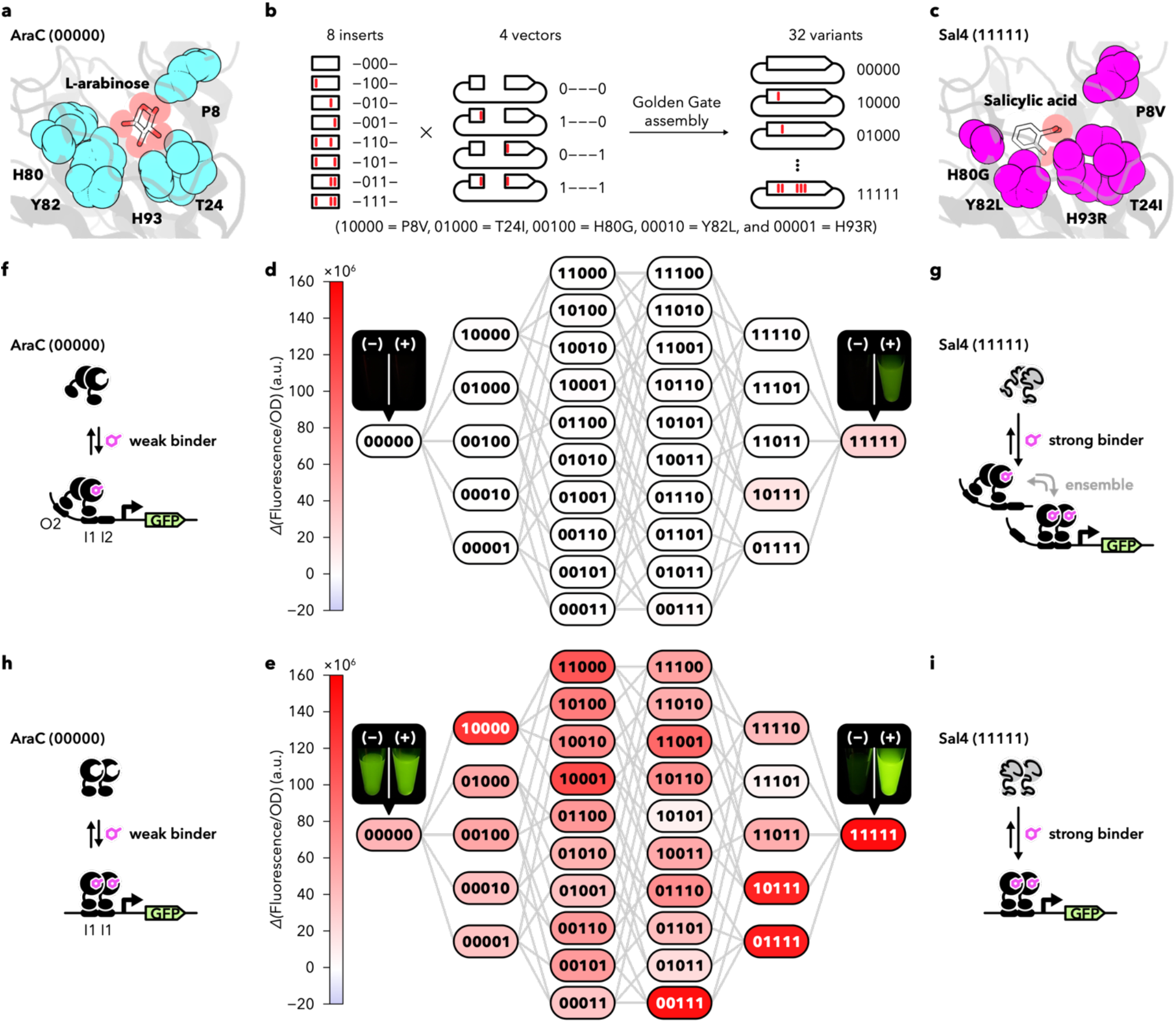
Comparison of the evolvability of salicylic acid response in allosteric and BIF modes. **a**, Residues targeted for generating Sal4. Five residues (cyan spheres) were subjected to mutagenesis. L-arabinose is shown as the stick and translucent sphere. The structure of AraC with L-arabinose was obtained from PDB ID 2arc. **b**, Construction of 32 AraC variants. Eight-insert DNA and four-vector DNA variants generated by PCR were subjected to Golden Gate assembly in all combinations to construct 32 AraC variant-expression plasmids. **c**, Binding pocket of Sal4 with five substitutions (shown as magenta spheres). Salicylic acid is shown as a stick and translucent sphere. The model structure of AraC with salicylic acid was generated using PyMOL and AutoDock vina. **d**,**e**, Responses of 32 AraC mutants to salicylic acid in allosteric and BIF modes, respectively. AraC and Sal4 alleles are indicated by 0 and 1, respectively. The mean of the *Δ*(Fluorescence/OD) of three parallel experiments are shown. After dispensing 200 µL of cell culture into 1.5 mL microtubes, the fluorescent images were captured using a Gel Ninja. Quantitative fluorescence measurement and standard deviations are provided in **Supplementary Table 3**. An alternative representation is shown in **Supplementary** Fig. 3. Switching mechanisms of (**f**,**g**) AraC and Sal4 at P_BAD_ promoter and (**h**,**i**) at P_BAD7_ promoter.

In the allosteric mode (using P_BAD_ promoter activity as the readout), only two variants, Sal4 (11111) and a quadruple mutant (10111), exhibited a salicylic acid response (**Fig. 4d**), with a needle-like fitness landscape. Contrastingly, most of the same variants, including the single mutant (P8V, 10000), exhibited significant responses to salicylic acid in the BIF mode (using P_BAD7_ promoter activity as the readout) (**Fig. 4e)**. This demonstrates that although the mutations in the ligand-binding pocket of AraC facilitated novel binding mechanisms for salicylic acid while losing to bind L-arabinose (Extended Data Figs. 7b and 8), most of them did not exhibit allosteric activation of AraC. Thus, the success rate of developing new sensor functions can be dramatically increased by adopting binding-induced stabilisation.

Interestingly, Sal4, which was originally isolated during the screening for salicylic acid-induced P_BAD_ activation, exhibited a higher signal output and stringency with P_BAD7_ activation (BIF mode). Therefore, the five mutations necessary for salicylic acid response (in Sal4) could significantly destabilise AraC, and possibly make it a salicylic acid-induced folder (**Figs. 4h** and i). If salicylic acid-induced folding stabilisation applies equally to the active (binding to the I1–I2 sites) and inactive (binding to the I1–O2 sites) forms (**Figs. 4f and g**), Sal4 still needs to evolve to acquire an optimal allosteric response to salicylic acid.

### Cooperation of BIF and allosteric modes

Antagonists competitively inhibit the allosteric response to ligands and agonists. When L-arabinose (AraC ligand) and D-fucose (AraC antagonist) are used as two input signals, the natural **AraC/P_BAD_** system is expected to behave as the NIMPLY gate; transcriptional activation is observed only in the presence of L-arabinose and absence of D-fucose (**Fig. 5a**). Contrastingly, AraC/P_BAD7_ system simply detects binding-induced stabilisation (**Fig. 1**) rather than the conformational changes associated with ligand binding. Hence, it does not distinguish between agonists and antagonists (**Fig. 2**) and is expected to behave as the OR gate that is activated in the presence of either L-arabinose or D-fucose or both. Indeed, using a plasmid with the RFP gene under the P_BAD7_ promoter, we confirmed its OR gate behaviour (**Fig. 5b**). Therefore, the same AraC variant can control the expression of two different promoters using completely different logic operations.

**Fig. 5.**
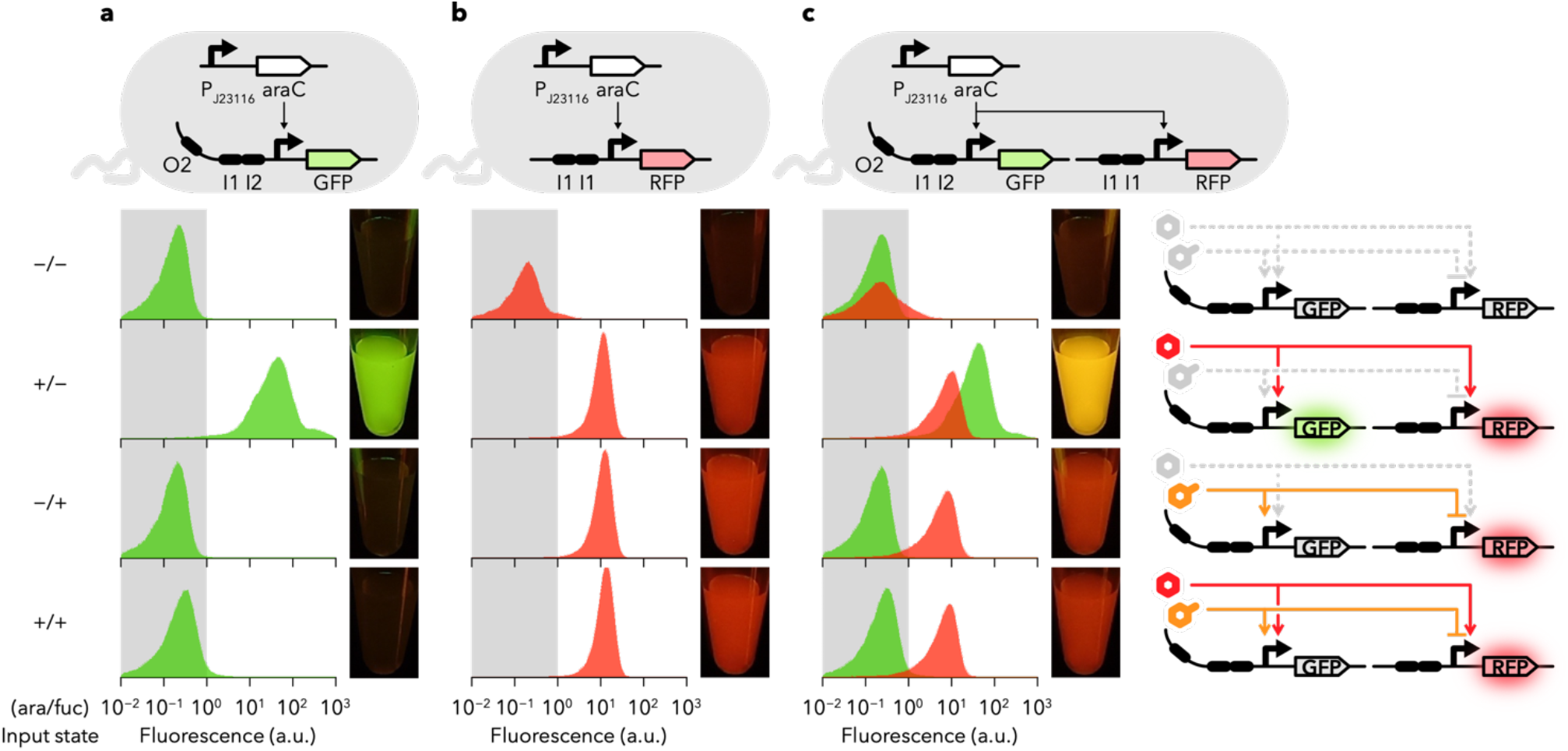
A single AraC variant behaves as a ‘selective NIMPY-OR converter’. **a**,**b**,**c**, Cell populations and culture images harbouring AraC_G1P1G4_/P_BAD_, AraC_G1P1G4_/P_BAD7_ and AraC_G1P1G4_/P_BAD_/P_BAD7_ systems, respectively. All logic functions were tested in the presence or absence of 1 mM L-arabinose or D-fucose. From top to bottom: no addition, L-arabinose only, D-fucose only and both. Green and red histograms indicate green and red fluorescence, respectively. Grey areas indicate no significant fluorescence, i.e. “off” state. Representative cell populations from four independent experiments are shown. Further details are provided in **Extended Data Fig. 9**.

We found that genes controlled by P_BAD_ (GFP) and P_BAD7_ (RFP) can coexist in the same cell and both promoters can be activated simultaneously and independently by the same AraC variant acting as a selective NIMPLY-OR converter (**Fig. 5c**). To our knowledge, there are no reports of a dual-output logic gate employing a single transcription factor.

### Integration of allosteric and BIF modes

We found that most of the AraC mutants activated the P_BAD7_ promoter at lower concentrations of L-arabinose than the P_BAD_ promoter (**Extended Data Fig. 2**). In the case of the AraC mutant G1P2G7, P_BAD7_ response was five times more sensitive to arabinose (EC_50_ = 21 µM) than that of P_BAD_ (EC_50_ = 99 µM) (**Fig. 6a**), possibly due to the difference in the operator sites that AraC acts on. In the P_BAD_ promoter, the binding of the AraC dimer to the I1-O2 sites is 18 times stronger than to the I1-I2 sites^18^. The apo form of AraC is selectively stabilised by O2 binding, thereby decreasing the extent of ligand (arabinose)-induced stabilisation. Hence, arabinose-induced conformational changes and the shifting to I1-I2 sites of AraC are shifted toward a higher concentration of arabinose, which does not represent the equilibrium between L-arabinose and AraC in the absence of operators. Contrastingly, the P_BAD7_ site lacks the O2 site, therefore, its arabinose response might directly reflect the equilibrium between L-arabinose and AraC.

**Fig. 6.**
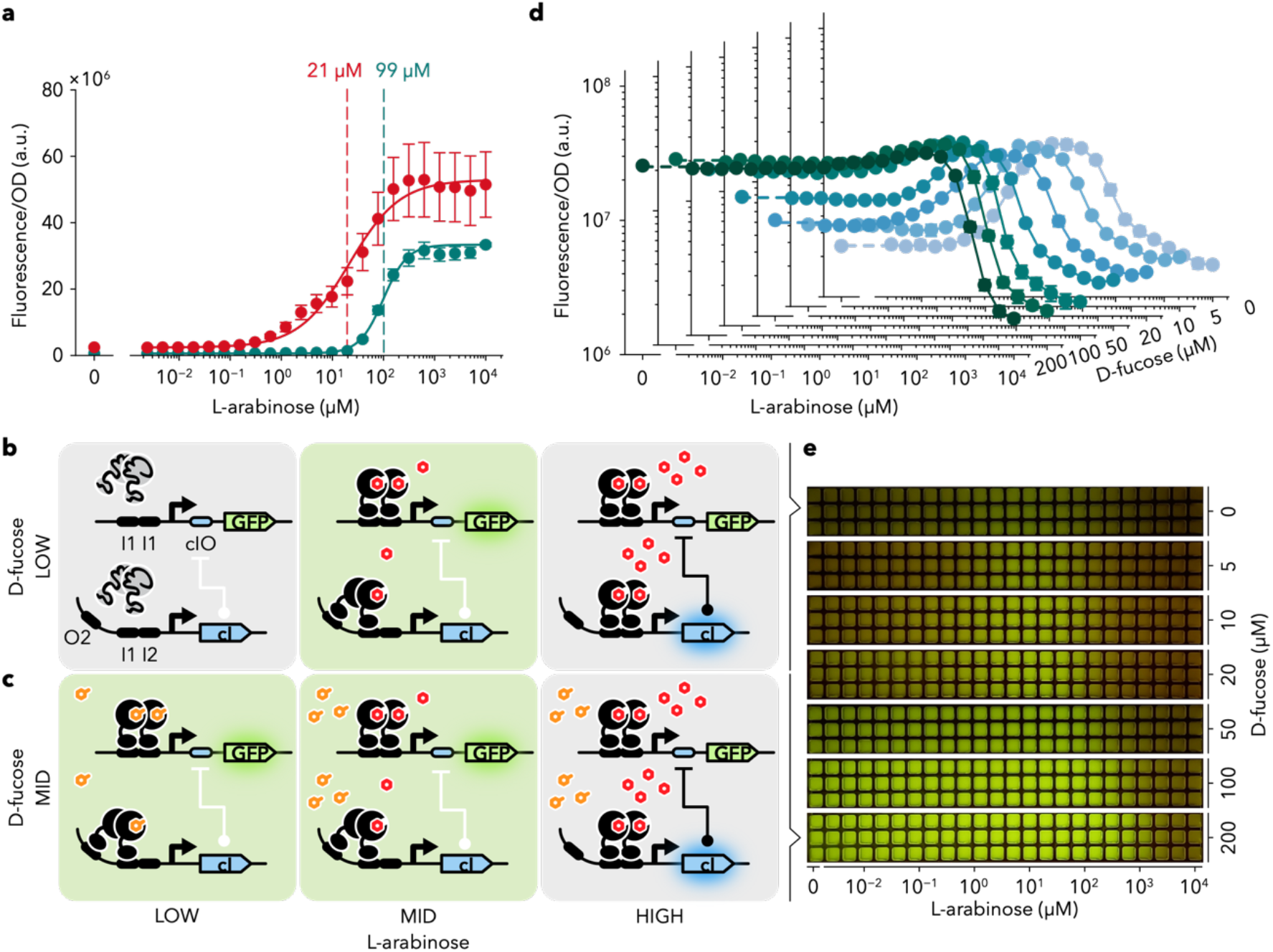
Integration of BIF and inverted allosteric output yield tuneable band-pass filter. **a**, Dose response of AraC_G1P2G7_ to L-arabinose in allosteric (green) and BIF (red) modes. The EC_50_ values for both modes were determined by the Hill equation. **b**, Design of band-pass filter. At low L-arabinose concentrations, both promoters are inactive due **Fig.** to the low stability of AraC. At moderate arabinose concentrations, stabilised AraC activates transcription at the P_BAD7/cIO_ promoter, while, at high arabinose concentrations, it also activates the P_BAD_ promoter to express the cI protein, which strongly represses transcription at the P_BAD7/cIO_ promoter. **c**, Design of transition to a low-pass filter: The presence of D-fucose further stabilises AraC and slightly lowers the switching threshold of the arabinose high-pass filter. D-fucose antagonises the P_BAD_ (allosteric) gene expression, thereby elevating the switching threshold of the low-pass filter. Consequently, the bandwidth of this BPF circuit is largely expanded. **d**, *E. coli* performances that installed the variable filter circuit. **e**, Images of cell functions. After dispensing 100 µL of cell cultures used in **c** into 384-well shallow plates, fluorescent images were captured using a Gel Ninja. In **a** and **d**, the data corresponds to the mean ± s.d. of three biological replicates.

AraC mutants respond to each promoter, which can be independently regulated within the same cell (**Fig. 5**), with different detection limits. Band-pass filter (BPF) is a circuit used to monitor whether a certain substance exists within the correct concentration range. Using the synthesis of morphogens as the output, BPF can be applied for pattern formation at a macroscopic scale. BPF can be constructed by genetically integrating a low- and high-pass filter (LPF and HPF) with different detection limits for the input molecule^40^. The mechanism of BPFs has been shown to include the simultaneous detection of various concentrations of input molecules by receiver proteins and the expression of several regulatory proteins, exploiting the differences in affinity of these regulators to DNA. Here, at least two or more transcription factors and/or polymerases are required with the receiver protein^41^.

We thought it was feasible to integrate the outputs from the P_BAD_ and P_BAD7_ promoters to develop a BPF with fewer components (**Fig. 6b**). Therefore, to integrate the outputs of both P_BAD_ and P_BAD7_ promoters, the lambda cI repressor gene was placed under the P_BAD_ promoter and its binding site, cIO, was inserted into the core region of the P_BAD7_ promoter (called the P_BAD7/cIO_ promoter, Extended Data Table 1) to invert the output from P_BAD_ promoter.

As expected, the resultant genetic circuit behaved as a BPF (**Figs. 6d and e**). Compared to the previously reported BPFs, this BPF circuit is unique as a single sensor protein, AraC, plays a dual role of an LPF and an HPF. The most remarkable feature of this BPF is that it is constructed by integrating the BIF and allosteric modes of AraC. Here, D-fucose agonises the BIF component, but antagonises the allosteric component (**Fig. 6c**). Consequently, this BPF circuit continuously transitions from BPF to LPF with increasing concentrations of D-fucose (**Figs. 6d and e**).

## Discussion

Sensors operating on the BIF principle are expected to have a high probability to invent new sensor functions, as they can act as a molecular switch only by fabricating interface that binds to the target molecules. To test this hypothesis, it is necessary to allow BIF-mode and allosteric mode to coexist in a single sensor protein, enabling the direct comparison of the frequency of new functions in the same mutant library of the protein. For this purpose, we employed a promoter configuration (P_BAD7_) in which AraC effectively acts as a super-activator (non-switch). Upon de-stabilization by mutation, AraC rapidly acquired BIF-mode while maintaining its original function as an allosteric sensor (P_BAD_). This enabled us, for the first time, an experimental comparison of the evolutionary capacities (capacities to invent new sensory functions) of AraC operating in BIF-mode and in allosteric modes. It revealed that AraC evolves new target response in remarkably high frequency, at least for the three different targets, D-fucose (natural antagonist of AraC; **Fig. 2**), D-galactose (non-binder; **Fig. 3**), and salicylic acid (irrelevant compound; **Fig. 4**).

Originally, BIF^1–4^ was proposed as an attractive approach that enables the rapid development of molecular switches and sensors without the necessity for laborious task of designing binding-induced allosteric modulation. Proteins are only marginally stable in nature, and folding energy can be cancelled by introducing a handful of mutations. As a phenomenon, BIF has been recognised for decades: many proteins are known to be better purified or crystallised as complexes with their ligands, substrates, or inhibitors^42–45^, and variants of sensory proteins behaving as BIFs have been described in mutational studies^46–49^. In our recent work, almost 20% of the non-switching variants of quorum sensor protein LuxR turned into stringent switchers upon random mutagenesis^4^. This surprisingly high frequency of emergence of switchers was also observed with AraC (**Fig. 1**). Thus, any protein can be readily transformed into a BIF through random mutation, highly accessible resource during evolution.

The newly added sensory function of AraC (non-allosteric regulation of P_BAD7_) is not only fully compatible with, but also qualitatively different from the original function (allosteric regulation of P_BAD_). As far as we known, this is the first report on a single-protein machinery that can operate two distinct regulatory behaviours. First, the sensors operating in BIF-mode are the visualizers of the stabilisation upon target binding, and therefore are unique in that they do not distinguish between agonists and antagonists. Secondly, sensitivity of BIF-sensors can be freely modulated (**Extended Data Fig. 2**), by tuning either of the target affinity by mutation or by changing operator configurations. Thus differentiated two sensory functions can be integrated into unique circuitry behaviours like a selective NIMPLY-OR converter (**Fig. 5**) and band pass filter with bandwidth tuning function (**Fig. 6**). Proteomic analysis has unveiled that proteins interact with a surprising number of metabolites and other proteins, which have non-negligible effects on their stability^50,51^. Binding-induced stabilisation enables all such interactions to be active participants in the decision making of the behaviours of physiological network.

Mutation drives protein evolution whilst simultaneously being the primary evolutionary constraint. Protein engineers are striving to obtain adaptive mutations that bestow new functions, whilst avoiding the destabilising effects of mutations^15–21^. Chaperone over-expression^52^ and stabilising mutations^53,54^ have been proven to be highly effective in mitigating this destabilisation effect. This study highlights a unique scenario in which mutations’ destabilising impact may accelerate the evolution of protein function. Well-evolved allosteric enzymes frequently acquire BIF properties due to mutational instability. This makes the newly invented novel target binding selectable trait. Given that binding-induced folders are frequently emerged from complex allosteric protein machinery (**Fig. 1**), it is tempting to speculate that the new binding properties are generated first in the non-allosteric mode, leading to sensors that exhibit binding-induced conformational changes and finally resulting in a mature allosteric sensor. These evolutionary intermediates may subsequently be upgraded to full-fledged allosteric sensors, driven by the evolutionary requirement for selectivity and/or economic demands in protein biogenesis. Exploring through BIF will also enable the development of novel biosensors tailored for the ever-increasing repertoire of molecules, either discovered in nature or created by chemists.

## Supporting information

supplementaldata-kimura-etal

## Methods

### Bacterial strains, media and growth conditions

*Escherichia coli* strains, JW0063 (ref.^55^), with an eliminated kanamycin-resistance cassette (**Supplementary** Fig. 4) and XL10-Gold (Agilent Technology, Inc., Santa Clara, CA), were used for cloning and library construction. Genotypes are provided in Supplementary **Table 4**. JW0063 harbouring P_BAD_-sfgfp-HSVtk-aph or P_BAD7_-sfgfp-HSVtk-aph was used as the reporter/selector strain for the directed evolution of AraC. For all the experiments, except the salicylic acid one, *E. coli* strains were grown at 37°C using appropriate antibiotics at the following concentrations in LB liquid medium (2% w/v) Lennox LB; Nacalai Tesque, Kyoto, Japan) or LB-agar plates (2% w/v) Lennox LB; Nacalai Tesque, 1.5% (w/v) agar; Nacalai Tesque), with 100 µg/mL of ampicillin (Sigma-Aldrich, Inc., St. Lous, MO), 30 µg/mL of chloramphenicol (Nacalai Tesque) and 30 µg/mL streptomycin (Nacalai Tesque). For the main culture in the salicylic acid experiment, the LB-TES medium was buffered with 50 mM TES (Nacalai Tesque) and adjusted to pH 7 using NaOH. The kanamycin (Sigma-Aldrich) stock solution (30 mg/mL) for the ON-state selection was prepared by dissolving appropriate amounts of kanamycin in deionised water and filter-sterilising it through a 0.2 µm cellulose acetate filter (MN Steriliser CA, Macherey-Nagel GmbH & Co. KG, Düren, Germany). The L-arabinose (Tokyo Chemical Industry Co., Ltd., Tokyo, Japan) and D-fucose (Tokyo Chemical Industry Co., Ltd) stock solutions (both 1 M) were prepared by dissolving appropriate amounts of the compounds in deionised water and filter-sterilising through a 0.2 µm cellulose acetate filter. The salicylic acid (Nacalai Tesque) stock solution (500 mM) was prepared by dissolving appropriate amounts of the compound in ethanol. Plasmid list, primer list and plasmid information are provided in **Supplementary Table 5, 6** and **Supplementary Note 1**, respectively.

### Library construction

#### Whole-gene mutagenesis

High-fidelity PCR was conducted using KOD DNA polymerase (TOYOBO, Osaka, Japan) to amplify the *araC* region or the mutants on the pET-based vector. The resulting fragment was subjected to error-prone PCR under the following conditions: 5 U of Taq DNA polymerase (New England Biolabs, MA, USA), 200 μM of each deoxynucleoside triphosphate, 2 mM of MgCl_2_ and 50 μM of MnCl_2_. The amplification factor was approximately 10^3^. The PCR product was digested at the *Nco*I and *Bam*HI sites and ligated into a pET-based vector. The ligation mixture was transformed into JW0063 by electroporation. DNA from the transformants was extracted via miniprep to yield the library plasmid (library size is approximately 10^6^).

#### Site-saturated mutagenesis

Site-saturation mutagenesis was induced for F15, M42 and I46 using ExSite PCR with the primers containing the NNK sequence (N is an equimolar mixture of dATP, dCTP, dGTP and dTTP; K is an equimolar mixture of dGTP and dTTP) at the targeted sites. After transforming the resultant plasmids into XL10-Gold, the transformants were plated on LB agar plates and grown overnight. Approximately 10^3^ colonies from each plate were scraped and pooled and their plasmid DNA was extracted by miniprep.

#### Gene expression ahnalysis using fluorescent proteins as reporters

For quantitative assays, JW0063 harbouring P_BAD_-sfgfp-HSVtk-aph or P_BAD7_-sfgfp-HSVtk-aph and plasmids encoding AraC mutant genes were first grown overnight from single colonies and then 1% cultures were inoculated into 400 µL of LB medium containing appropriate antibiotics and L-arabinose and/or D-fucose in 96-deep well plates. These cultures were shaken at 37°C for 12 h. Then, 20 µL cultures were diluted 10-fold with saline (0.9% (w/v) NaCl; Nacalai Tesque) in 96 shallow-well plates. Cell densities were measured using FilterMax F5 (Molecular Devices, San Jose, CA) at 595 nm. The GFP fluorescence (excitation at 485 nm and emission at 535 nm) and RFP fluorescence (excitation at 585 nm and emission at 625 nm) were measured using the FilterMax F5. Fluorescence values were normalised using OD_595_. Sensor function was defined as the difference in fluorescence intensity per OD between the two conditions. This was calculated using the following equation:

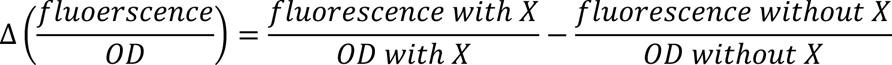

where X indicates target molecules, i.e. 1 mM L-arabinose, 1 mM D-fucose, 10 mM D-galactose and 5 mM salicylic acid.

Fluorescence characterisation with flow cytometry was performed on a MACS Quant VYB (Miltenyi Biotech, Bergisch-Gladbach, Germany). The cell cultures grown in the presence or absence of 1 mM L-arabinose and/or 1 mM D-fucose were diluted 1:50 with 200 µL of saline in a 96 shallow-well plate. We counted and measured 5 x 10^4^ cells using an FSC voltage of 320 V, an SSC voltage of 230 V, a B1 laser (excitation at 488 nm and emission at 525/50 nm) voltage of 420 V and an Y2 laser (excitation at 561 nm and emission at 615/20 nm) voltage of 400 V. The data were analysed using a MACS Quant analyser (Miltenyi Biotech, Bergisch-Gladbach, Germany).

### Prediction of the stability change using FoldX

The model structure of full-length AraC (ID: AF-P0A9E0-F1-model_v2) predicted by AlphaFold2 and published in UniProt was used. To eliminate unfavourable torsion angles, van der Waals’ clashes or total energy, the side chains of the model structure were rearranged using the RepairPDB command in FoldX to generate a stabilised structure. The free energy change between the wild-type and mutant (ΔΔG) was predicted using the BuildModel command with the following configuration: ionStrength = 0.05, pH = 7, temperature = 298, vdwDesign = 2, moveNeighbours = true and a number of runs = 3. The mean of the three runs was used in the analysis.

## Data and code availability

Not applicable.

## Acknowledgements

This project is a part of the outcome of research performed under a JSPS fellowship for young scientists (17J08108), a Waseda University Grant for Special Research Projects (2022C-465) and a Nissin Sugar Foundation.

## Author contributions

Y.K. and D.U. conceived the project and designed the experiments; Y.K. performed all experiments with assistance from S.K.-N. and D.U.; Y.K. and D.U. wrote the manuscript.

## Extended Data

**Extended Data Fig. 1.**
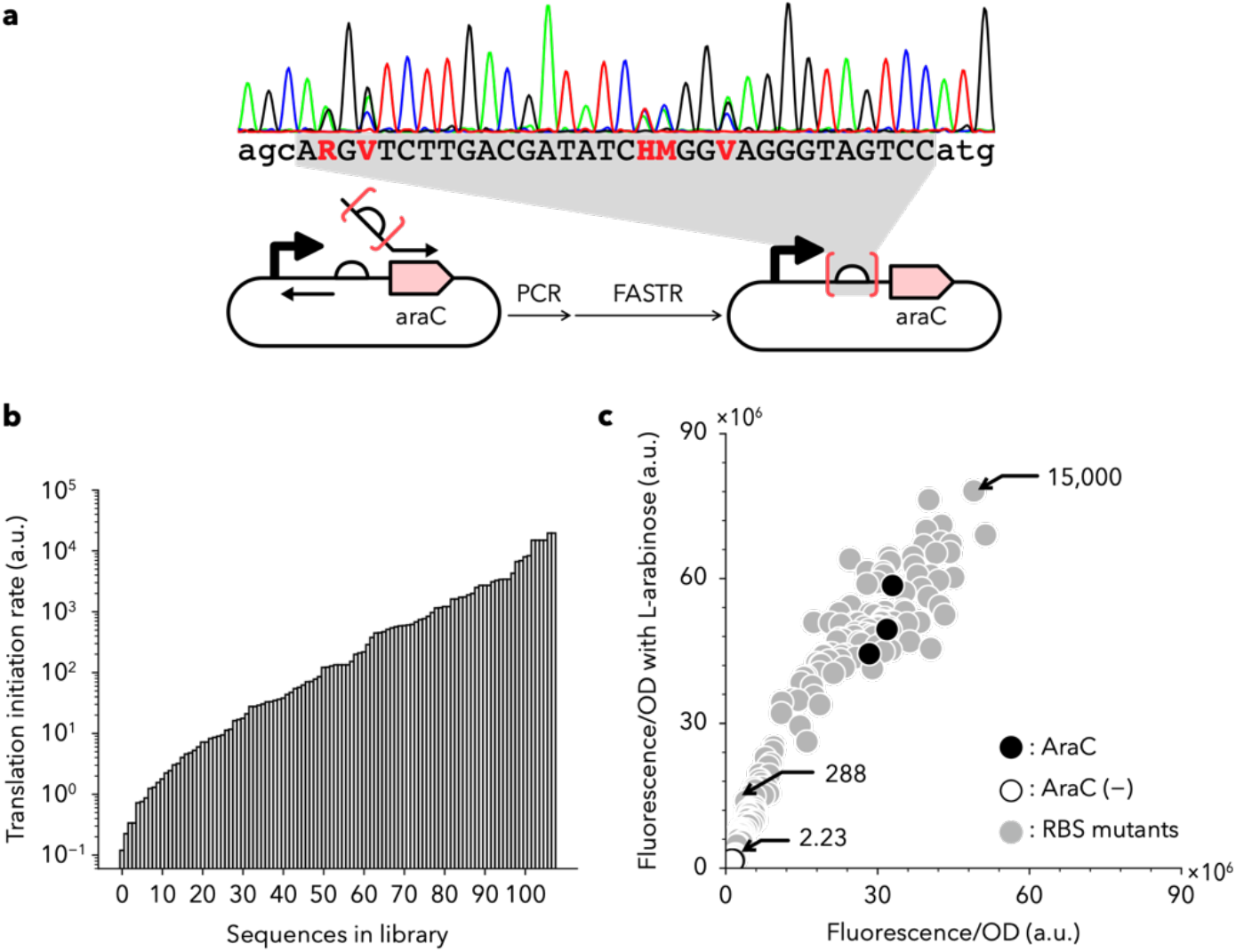
The effect of AraC expression level on the AraC/P_BAD7_ system. **a**, Construction of the ribosome binding site (RBS) library. The degenerated RBS was designed using the RBS Calculator and constructed using FASTR assembly. R, A or G; V, A, C or G; H, A, C or T; M, A or C. **b**, The distribution of RBS score. **c**, The switching function of the RBS variants with or without 1 mM L-arabinose. The values in the figure indicate the RBS score.

**Extended Data Fig. 2.**
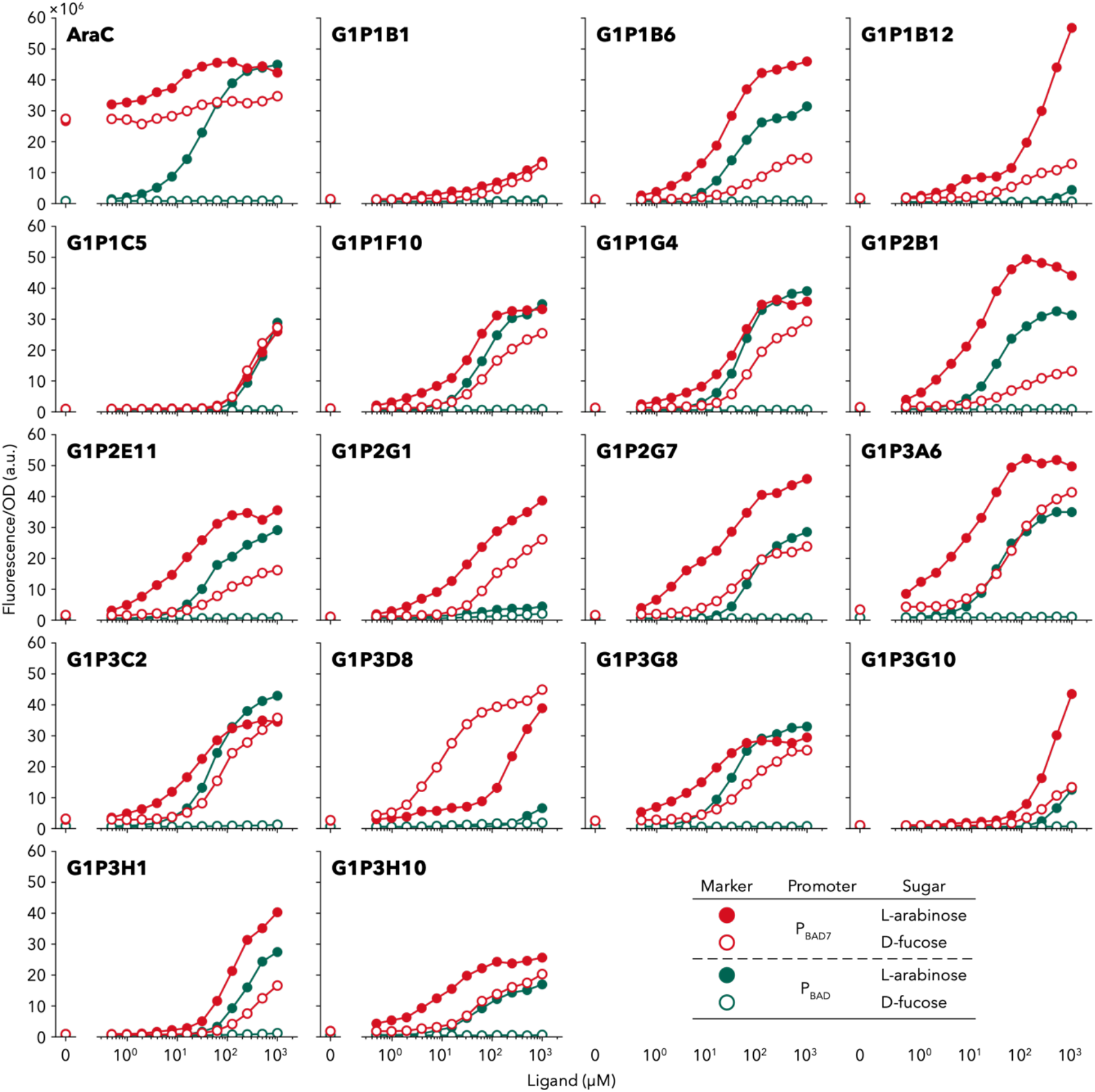
Dose response of AraC mutants at the P_BAD_ or P_BAD7_ promoters. The data points are connected by lines for better visualisation. These experiments were conducted as a single measurement.

**Extended Data Fig. 3.**
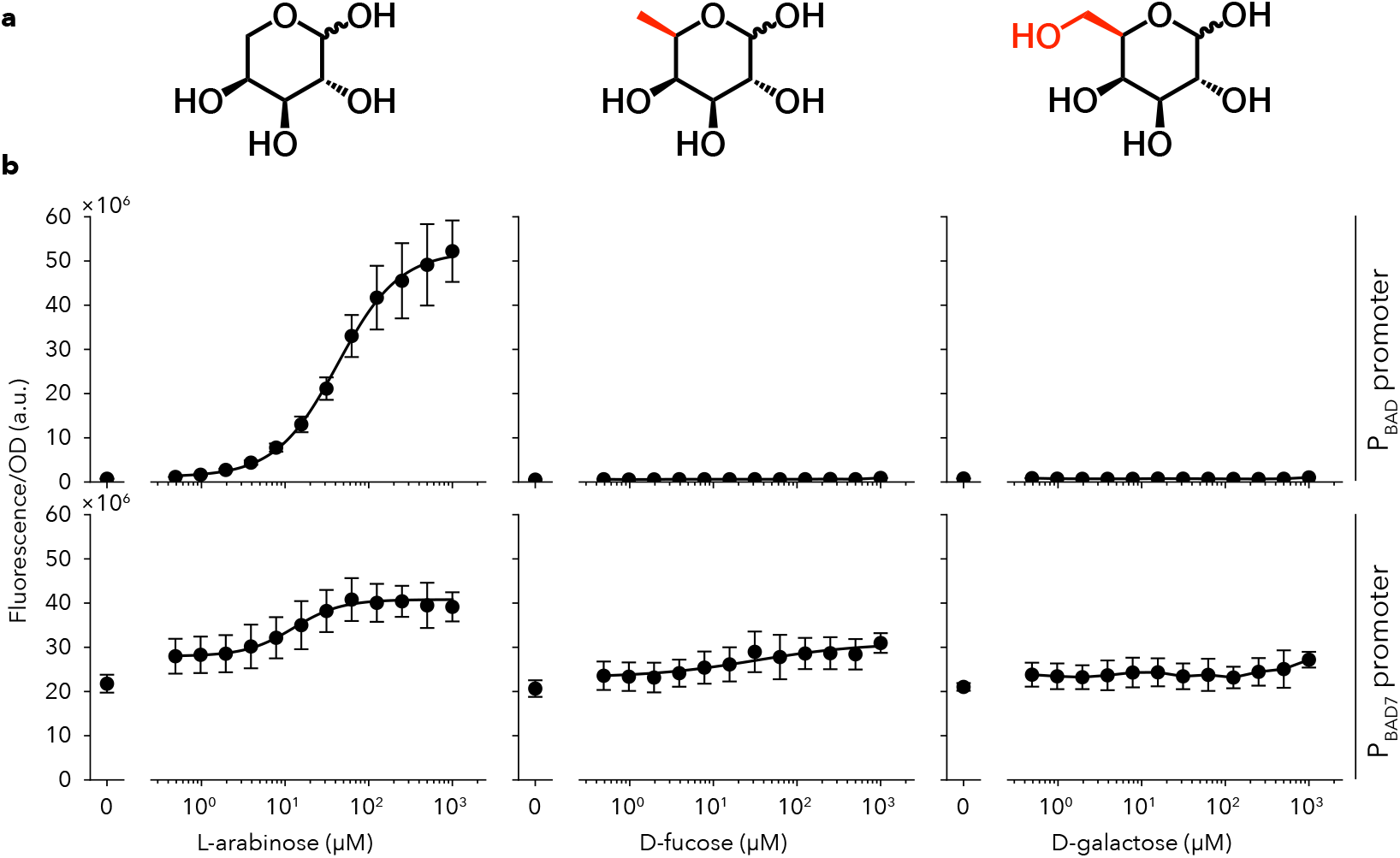
Target compounds and dose-response of wild-type AraC. **a**, Chemical structure of L-arabinose (native ligand), D-fucose and D-galactose. **b**, Transfer functions of wild-type AraC with L-arabinose, D-fucose and D-galactose at P_BAD_ and P_BAD7_ promoter. Each value corresponds to the mean ± s.d. of three independent experiments.

**Extended Data Fig. 4.**
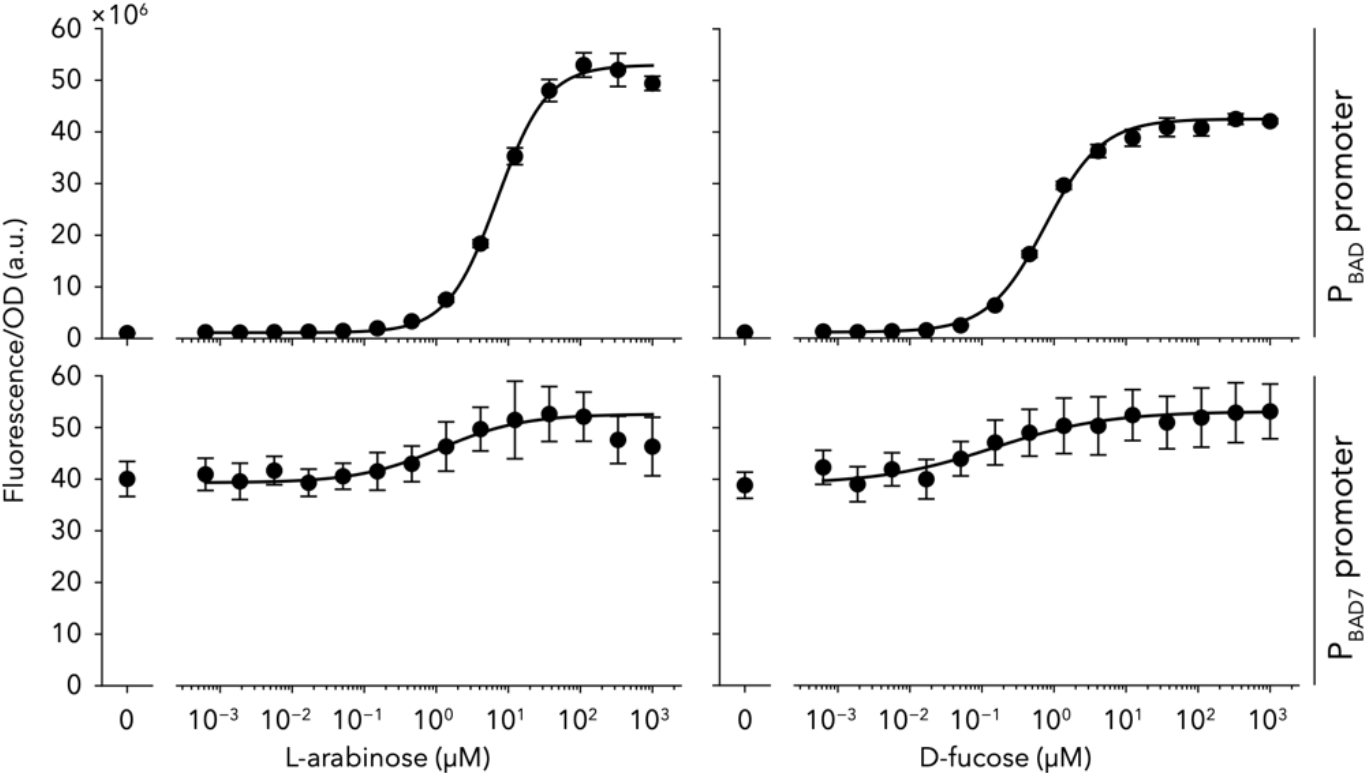
Effect of I46V on the dose-response to L-arabinose and D-fucose. Dose responses were obtained by fitting the data to the Hill equation. Each value corresponds to the mean ± s.d. of three biological replicates.

**Extended Data Fig. 5.**
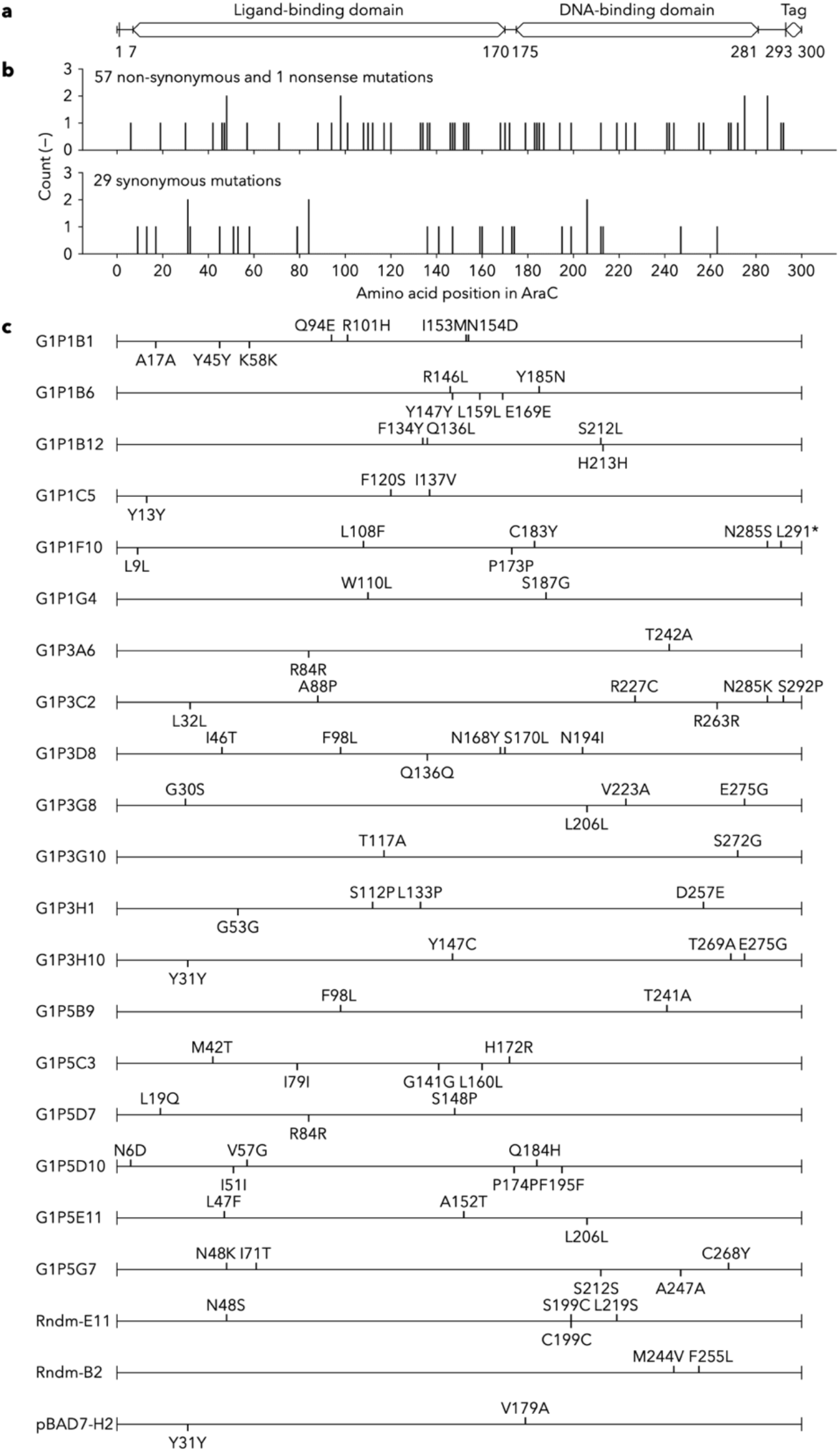
Distribution of mutations identified in selected D-fucose responders. **a**, Schematic diagram of *araC* gene. The ‘tag’ denotes a histidine-hexamer peptide. **b**, Histogram of the non-synonymous mutations and nonsense mutation (upper) and synonymous mutations (lower). The upper part of the bar graph shows 55 unique non-synonymous mutations, 2 duplicated non-synonymous mutations and 1 nonsense mutation. The non-synonymous mutations were analysed using FoldX and depicted in the upper panel of **Fig. 2e**. **c**. Mutation maps for each mutant were analysed using FoldX and depicted in the lower panel of **Fig. 2e**.

**Extended Data Fig. 6.**
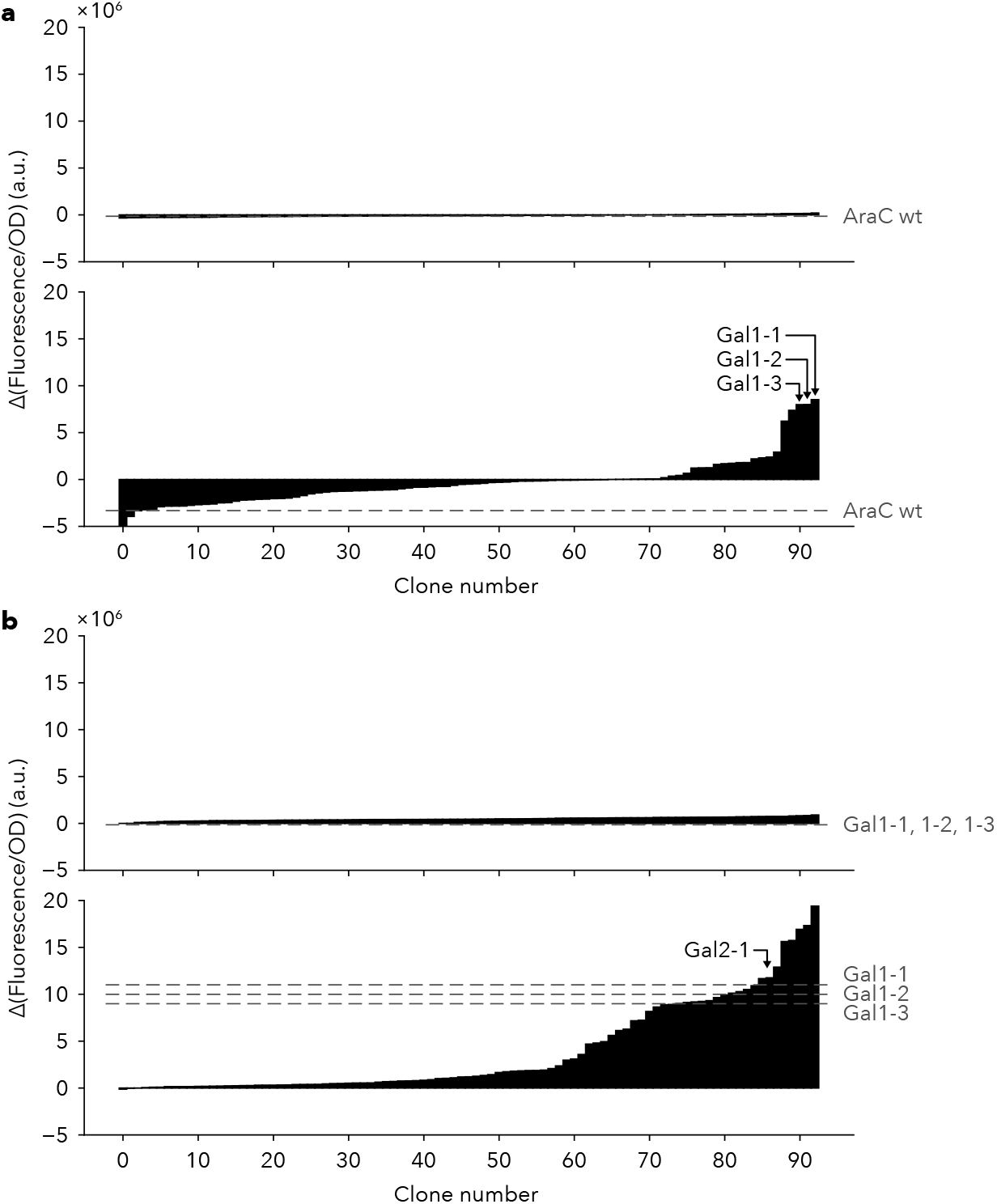
Directed evolution of AraC for D-galactose switch. **a**, Fitness landscapes of the 1st round for the allosteric (upper) and BIF (lower) modes. The function of 93 mutants randomly picked is represented as a filled bar and the function of wild-type AraC is shown as a grey dot line. **b**, Fitness landscapes of the 2nd round for the allosteric (upper) and BIF (lower) modes. The 2nd generation library was generated from a plasmid mixture of Gal1-1, Gal1-2 and Gal1-3 by error-prone PCR. The function of 93 mutants randomly picked is shown as a filled bar and the functions of Gal1-1, Gal1-2 and Gal1-3 are shown as grey dot lines. Note that Gal2-1 was elected because this mutant had a similar Δ(Fluorescence/OD) value to the parental mutants but was more stringent. In **a**,**b**, these experiments were conducted as a single measurement.

**Extended Data Fig. 7.**
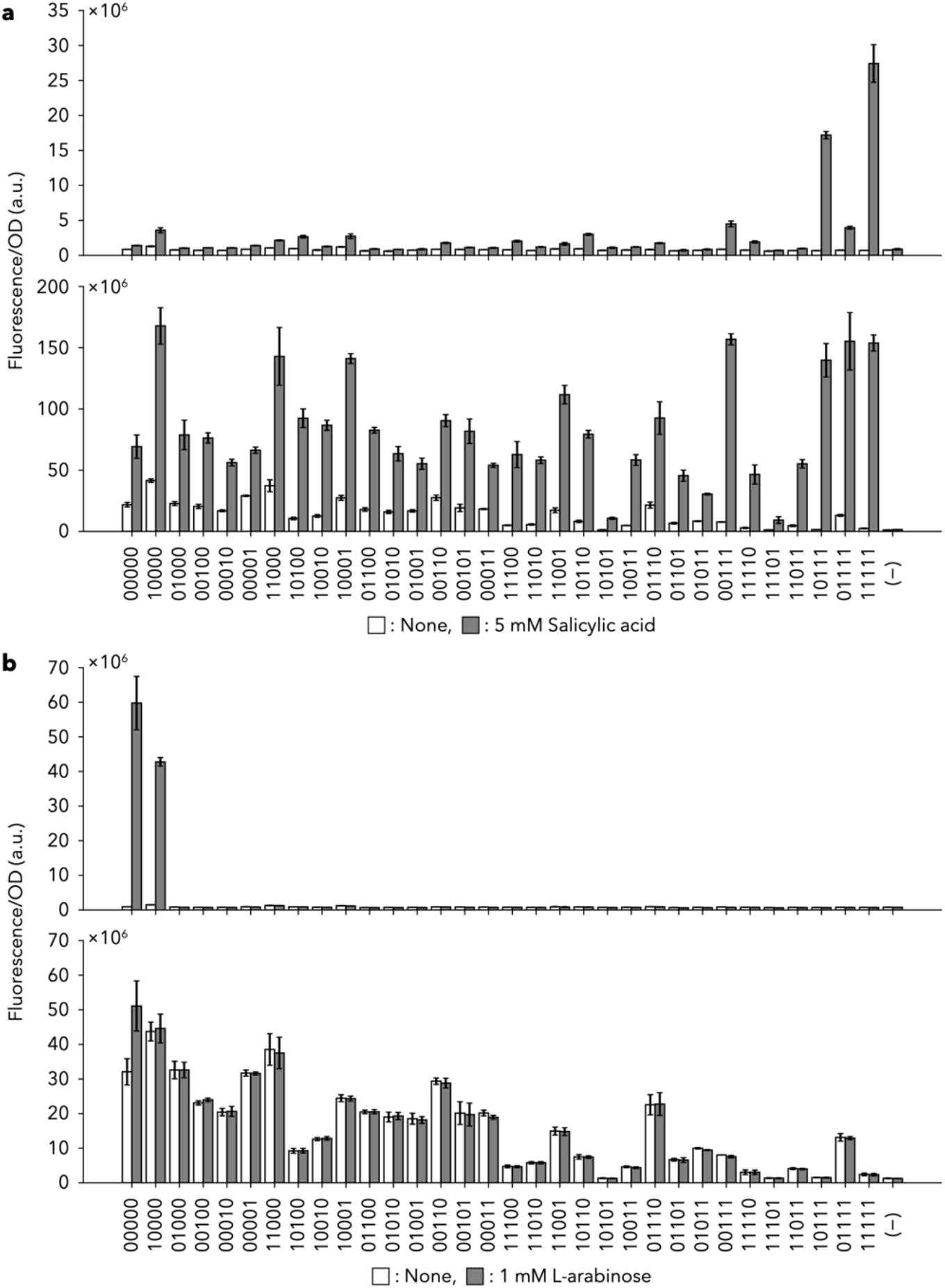
Salicylic acid (a) and L-arabinose (b) response of variants of Sal4, a mutant of AraC. **a**, The responses of the P_BAD_ (upper) and P_BAD7_ (lower) promoters, respectively, to 5 mM salicylic acid. **b**, The responses to the P_BAD_ (upper) and P_BAD7_ (lower) promoters, respectively, to 1 mM L-arabinose in LB-TES medium. In **a**,**b**, each value corresponds to the mean ± s.d. of 3 biological replicates. 10000, P8V; 01000, T24I; 00100, H80G; 00010, Y82L; 00001, H93R. (−) indicates data from *E. coli* that does not express AraC.

**Extended Data Fig. 8.**
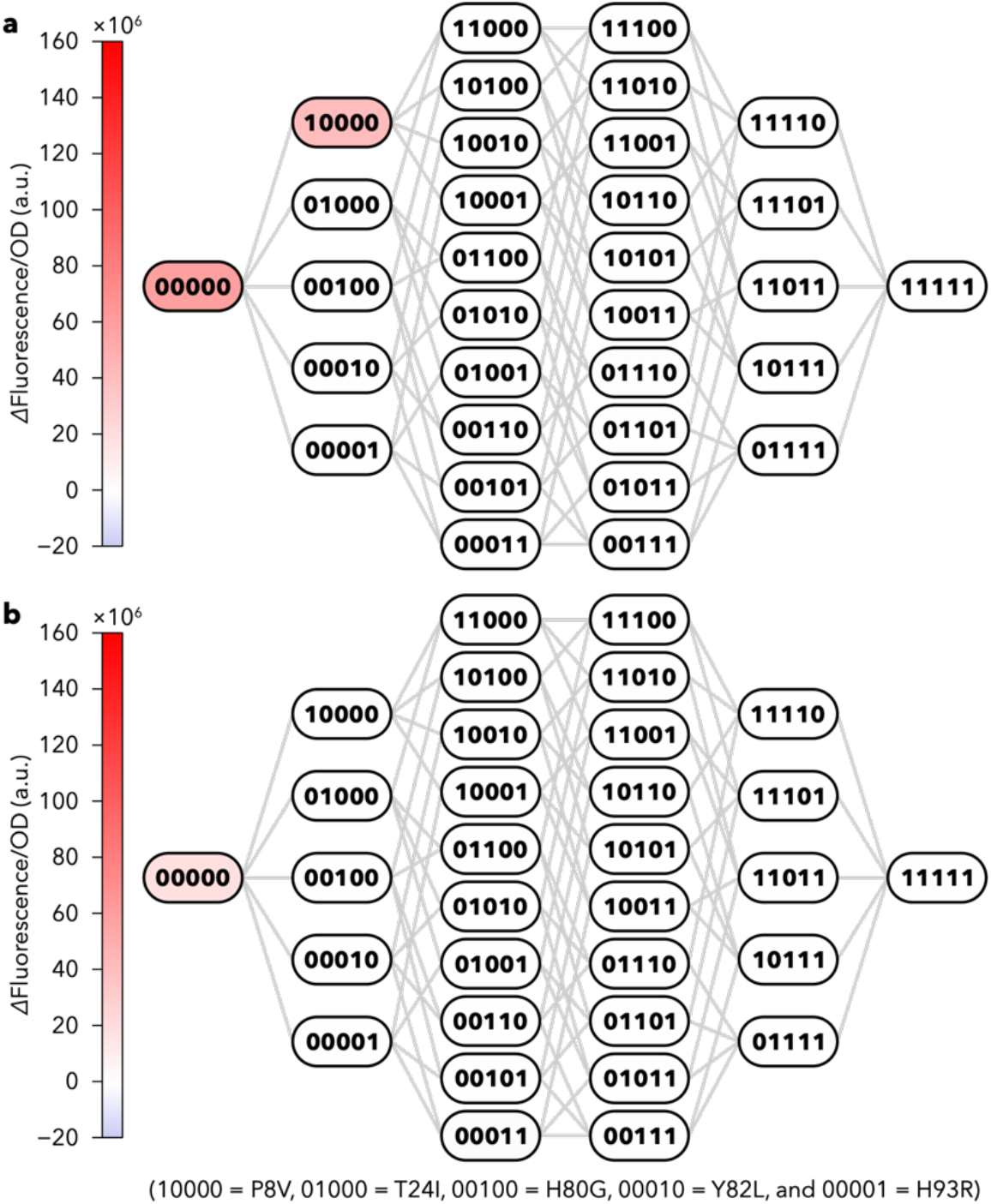
Fitness landscapes for L-arabinose response in the allosteric mode (a) and BIF mode (b). The differences between the averages of three parallel experiments in **Extended Data Fig. 7**b are shown.

**Extended Data Fig. 9.**
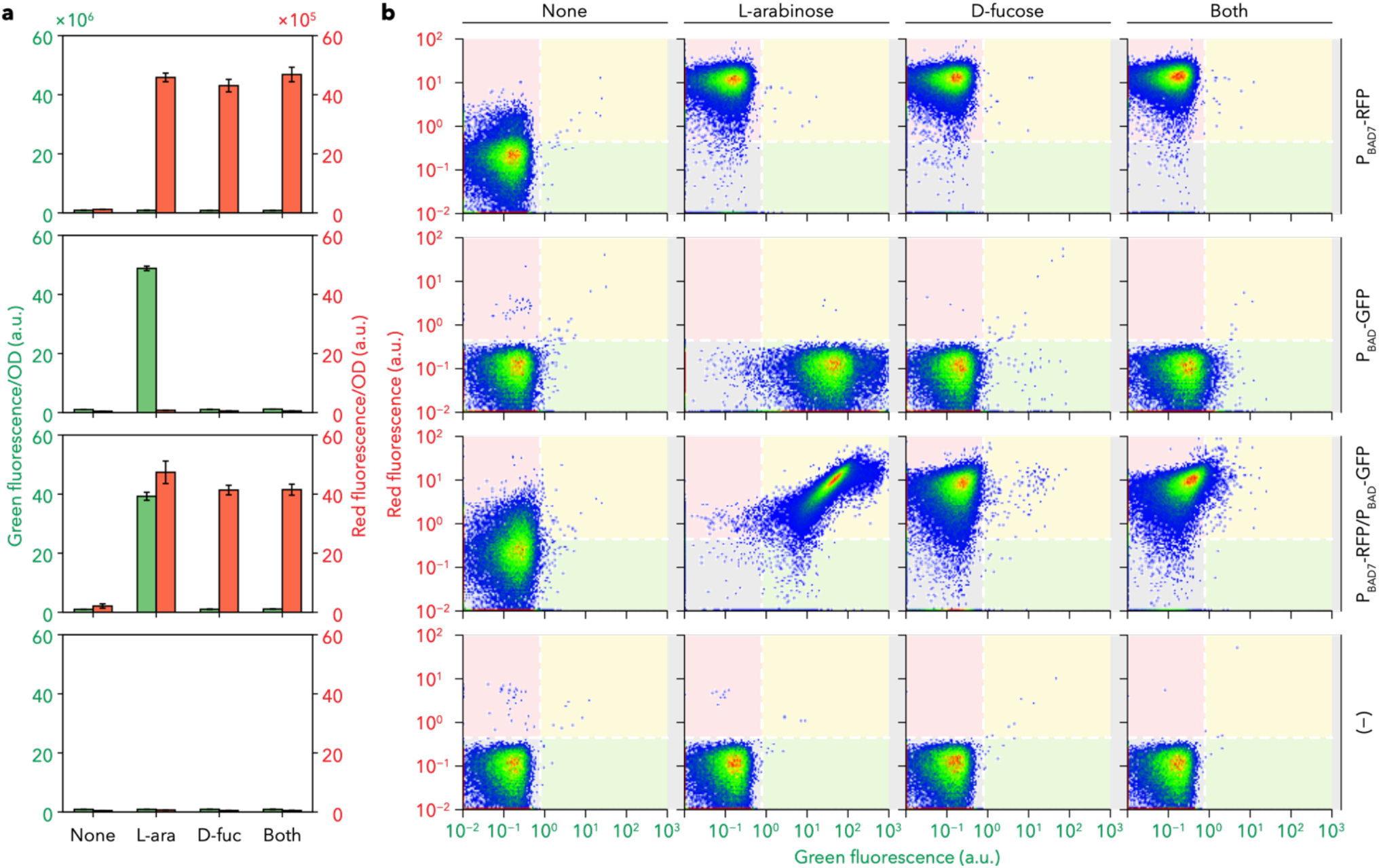
Construction of a ‘selective NIMPLY-OR converter’ that are simultaneously regulated by a sole AraC mutant in a single cell. **a**, Cell culture analysis. From top to bottom: logic function of cells harbouring P_BAD_-GFP, P_BAD7_-RFP, P_BAD_-GFP and P_BAD7_-RFP and no probe, respectively. Green and red filled bars indicate green and red fluorescence/OD, respectively. Data shown in bar graphs represent the mean ± s.d. from four experiments. **b**, Cell population analysis by flow cytometry. Representative cell populations from four independent experiments in **a** are shown.

**Extended Data Table 1.**
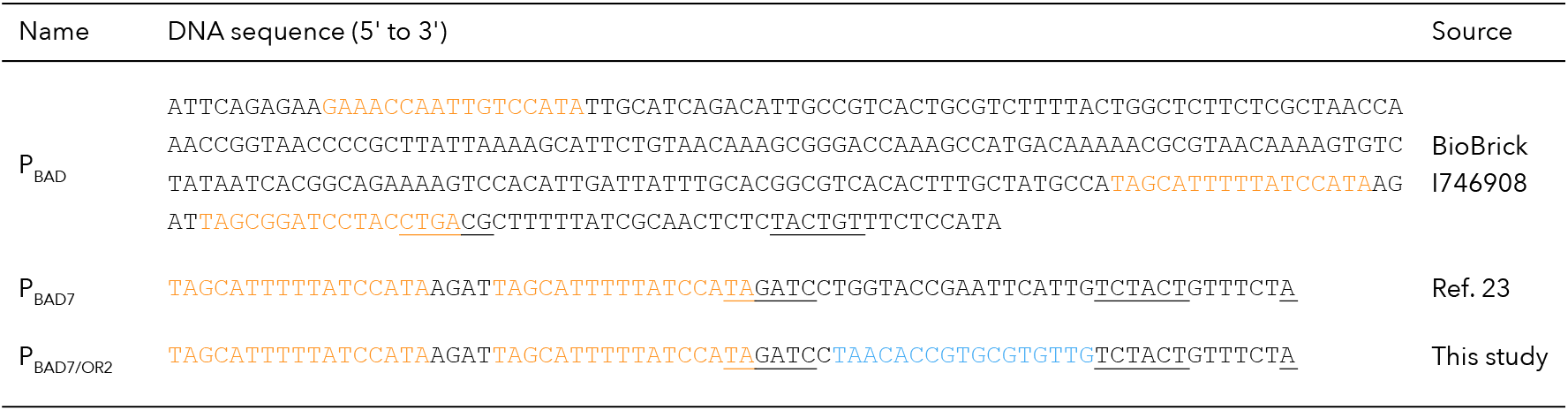
Promoter sequences regulated by transcription factors. −35 box, −10 box and +1 site are underlined. AraC-binding sites (O2, I1 and I2) and a lambda cI-binding site (cIO) are shown in orange and blue, respectively.

