## supplementaldata-kimura-etal for "Mutational destabilisation accelerates the evolution of novel sensory and network functions"

### Supplementary Figures

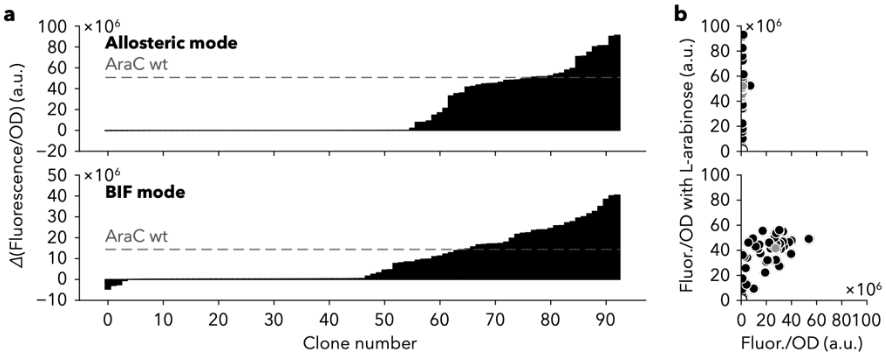


**Supplementary Fig. 1 | Comparing the fitness landscapes for the L-arabinose switch in the BIF and allosteric modes. a**, Fitness landscapes of AraC for the allosteric (upper panel) and non-allosteric (lower panel) modes. The function of 93 mutants randomly picked is represented as a filled bar and the function of wild-type AraC is shown as a grey dot line. Note that the data in the non-allosteric mode are also displayed in Fig. 1e. **b**, The data in **a** are replotted as a scatter plot with or without 1 mM L-arabinose. Black, mutants; grey, wild-type AraC; white, without AraC. These experiments were conducted as a single measurement.

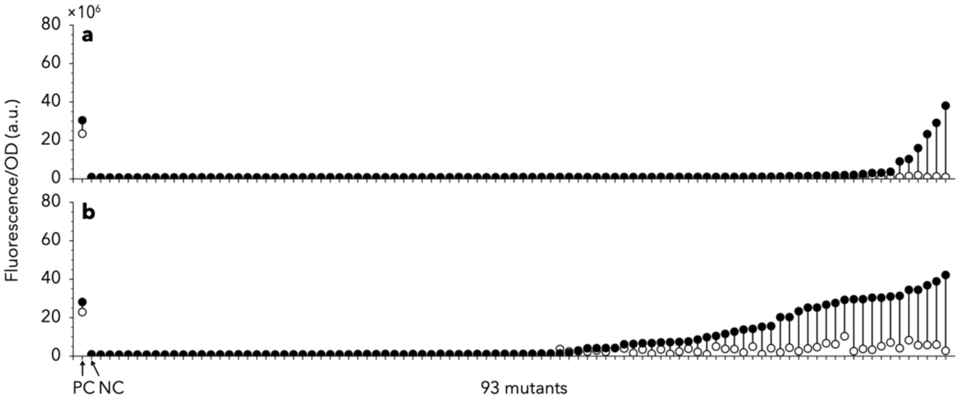


**Supplementary Fig. 2 | Directed evolution of AraC for D-fucose switch at the $P_{BAD7}$ promoter. a**,**b**, Distribution of mutants screened under off-state conditions only (**a**) and under off-state conditions after on-state screening with 1 mM D-fucose and 15 µg/mL kanamycin (**b**). Positive control (PC) and negative control (NC) refer to the cells harbouring the wild-type AraC plasmid and an empty plasmid, respectively. These experiments were conducted as a single measurement.

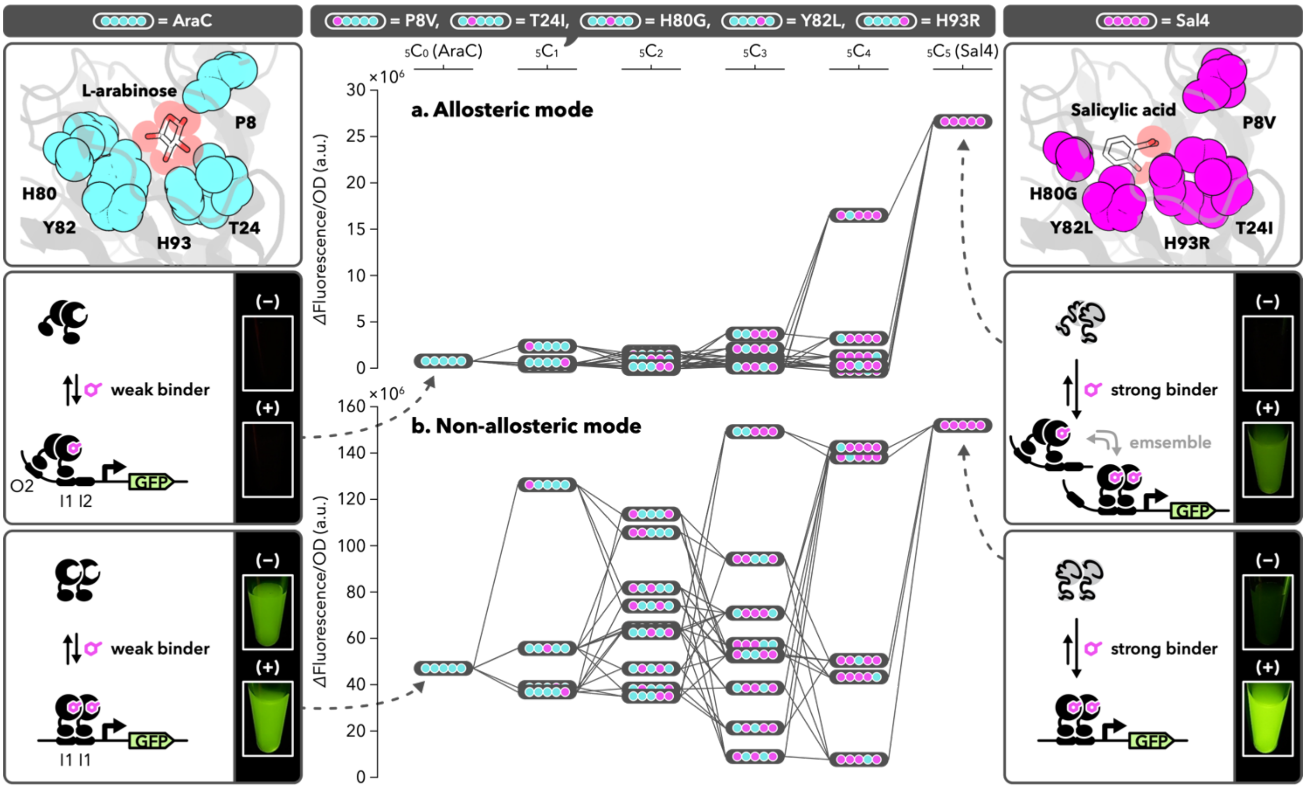


**Supplementary Fig. 3 | Fitness landscapes for salicylic acid response in the allosteric mode (a) and non-allosteric mode (b).** The differences between the averages of three parallel experiments in Extended Data Fig. 7a are shown. These data are also displayed in Fig. 4 but in a different format.

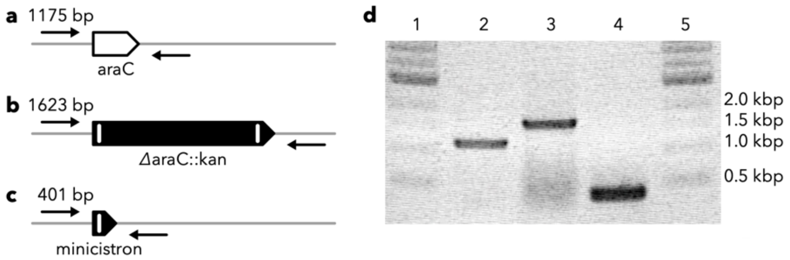


**Supplementary Fig. 4 | Construction of JW0063 deleted kanamycin-resistant gene. a,b,c,** Schematic diagrams of genomes in BW25113, JW0063 with a recombination cassette and the expected JW0063, respectively. Arrows represent the primer sets used in the genotype check in d. d, Gel electrophoresis of the PCR products amplified from each genome. Lane-1, NEB 1 kb DNA ladder; lane-2, BW25113; lane-3, JW0063 with a recombination cassette; lane-4, JW0063 without a recombination cassette; lane-5, NEB 1 kb DNA ladder.

### Supplementary Tables

#### Supplementary Table 1 | Mutations of AraC mutants.

| Name | Nucleotide mutations (putative amino acid mutations) | Screening condition |
|---|---|---|
| G1P1B1 | C51G (A17A), T135C (Y45Y), A174G (K58K), C280G (Q94E), G302A (R101H), A459G (I153M), A460G (N154D) | Run-1 |
| G1P1B6 | G437T (R146L), T441C (Y147Y), G477A (L159L), G507A (E169E), T553A (Y185N) | Run-1 |
| G1P1B12 | T401A (F134Y), A407T (Q136L), C635T (S212L), T639C (H213H) | Run-1 |
| G1P1C5 | C39T (Y13Y), T359C (F120S), A409G (I137V) | Run-1 |
| G1P1F10 | C25T (L9L), C322T (L108F), A519G (P173P), G548A (C183Y), A854G (N285S), T872A (L291*) | Run-1 |
| G1P1G4 | G329T (W110L), A559G (S187G) | Run-1 |
| G1P2B1 | G216T (L72F), T237C (I79I), G373A (A125T), A814G (S272G) | Run-2 |
| G1P2E11 | A573G (A191A), T794A (F265Y) | Run-2 |
| G1P2G1 | T6C (A2A), A47G (N16S), A53G (H18R), T66C (G22G), T210C (D70D), A694C (I232L), T784A (S262T) | Run-2 |
| G1P2G7 | T152C (I51T), T153C (T51T), A179G (Q60R), C203A (P68Q), C322T (L108F), C635T (S212L), G786A (S262S) | Run-2 |
| G1P3A6 | T252G (R84R), A724G (T242A) | Run-3 |
| G1P3C2 | C96T (L32L), G262C (A88P), C679T (R227C), A789G (R263R), T855A (N285K), T874C (S262P) | Run-3 |
| G1P3D8 | T173C (I46T), T292C (F98L), A408G (Q136Q), A502T (N168Y), C509T (S170L), A581T (N194I) | Run-3 |
| G1P3G8 | G88A (G30S), T616C (L206L), T668C (V223A), A824G (E275G) | Run-3 |
| G1P3G10 | A349G (T117A), A814G (S272G) | Run-3 |
| G1P3H1 | T159C (G53G), T334C (S112P), T398C (L133P), T771A (D257E) | Run-3 |
| G1P3H10 | T93C (Y31Y), A440G (Y147C), A805G (T269A), A824G (E275G) | Run-3 |
| G1P5B9 | T294A (F98L), A721G (T241A) | Run-4 |
| G1P5C3 | T125C (M42T), T237C (I79I), G423T (G141G), A480G (L160L), A515G (H172R) | Run-4 |
| G1P5D7 | T56A (L19Q), T252G (R84R), T442C (S148P) | Run-4 |
| G1P5D10 | A16G (N6D), T153A (I51I), T170G (V57G), G522A (P174P), G552C (Q184H), T585C (F195F) | Run-4 |
| G1P5E11 | C139T (L47F), G454A (A152T), T616C (L260L) | Run-4 |
| G1P5G7 | T144A (N48K), T212C (I71T), A636T (S212S), C741T (A247A), G803A (C268Y) | Run-4 |
| Rndm-E11 | A143G (N48S), A595T (S199C), C597T (C199C), T656C (L219S) | Run-4 |
| Rndm-B2 | A730G (M244V), T763C (F255L) | Run-4 |
| Rndm-H2 | T93C (Y31Y), T397C (V179A) | Run-4 |
| pBAD-A7 | A136G (I46V), T699C (S233S) | Run-5 |

Run-1, off-state screening ($P_{BAD7}$); Run-2, on-state selection with L-arabinose followed by off-state screening ($P_{BAD7}$); Run-3, on-state selection with D-fucose followed by off-state screening ($P_{BAD7}$); Run-4, random pick ($P_{BAD7}$); Run-5, random pick ($P_{BAD}$).

**Supplementary Table 2 | Mutations of D-galactose responsive mutants.**

| | Codon (residue) | | | | | |
|---|---|---|---|---|---|---|
| | 15 | 42 | 44 | 46 | 100 | 251 |
| AraC | TTT (F) | ATG (M) | GGT (G) | ATT (I) | CCG (P) | CGC (R) |
| Gal1-1 | TCG (S) | GAG (E) | GAT (D) | GAT (D) | CCG (P) | CGC (R) |
| Gal1-2 | TGG (W) | CCG (P) | GGT (G) | GAT (D) | CCG (P) | CGC (R) |
| Gal1-3 | TGG (W) | GAT (D) | GGT (G) | GAT (D) | CCG (P) | CGC (R) |
| Gal2-1 | TGG (W) | GAT (D) | GGT (G) | GAT (D) | CCA (P) | CTC (L) |

The mutation at the 44th codon in the Gal1-1 mutant was accidentally introduced in the 1st round. The mutations at the 100th and 251st codons were identified in the 2nd round evolution.

**Supplementary Table 3 | Sensor functions of AraC/Sal4 binary variants in each promoter.**

| | Residue | | | | | ΔFluorescence/OD/$10^6$ (a.u.) | | | |
|---|---|---|---|---|---|---|---|---|---|
| | | | | | | ± 5mM Salicylic acid | | ± 1 mM L-arabinose | |
| | 8 | 24 | 80 | 82 | 93 | $P_{BAD}$ promoter | $P_{BAD7}$ promoter | $P_{BAD}$ promoter | $P_{BAD7}$ promoter |
| 00000 | P | T | H | Y | H | 0.549 ± 0.0214 | 47.3 ± 7.86 | 58.8 ± 7.72 | 19.0 ± 3.76 |
| 10000 | V | T | H | Y | H | 2.30 ± 0.272 | 126 ± 15.8 | 41.3 ± 1.28 | 0.871 ± 5.40 |
| 01000 | P | I | H | Y | H | 0.267 ± 0.0283 | 56.0 ± 10.9 | −0.0820 ± 0.0569 | −0.00332 ± 1.20 |
| 00100 | P | T | G | Y | H | 0.366 ± 0.0556 | 55.8 ± 2.51 | −0.0674 ± 0.00813 | 0.895 ± 0.0976 |
| 00010 | P | T | H | L | H | 0.373 ± 0.0565 | 39.4 ± 2.17 | −0.0680 ± 0.0417 | 0.214 ± 0.872 |
| 00001 | P | T | H | Y | R | 0.518 ± 0.0542 | 37.2 ± 2.56 | −0.0825 ± 0.0394 | −0.103 ± 0.840 |
| 11000 | V | I | H | Y | H | 1.08 ± 0.0627 | 106 ± 19.2 | −0.0608 ± 0.0240 | −0.991 ± 1.43 |
| 10100 | V | T | G | Y | H | 1.70 ± 0.176 | 81.8 ± 6.69 | −0.0243 ± 0.00623 | 0.0515 ± 0.0544 |
| 10010 | V | T | H | L | H | 0.499 ± 0.0849 | 74.1 ± 3.06 | −0.0334 ± 0.00447 | 0.218 ± 0.214 |
| 10001 | V | T | H | Y | R | 1.53 ± 0.288 | 114 ± 2.80 | −0.101 ± 0.0228 | −0.130 ± 0.557 |
| 01100 | P | I | G | Y | H | 0.263 ± 0.114 | 64.8 ± 3.61 | −0.0281 ± 0.0175 | 0.0706 ± 0.0940 |
| 01010 | P | I | H | L | H | 0.252 ± 0.0823 | 47.5 ± 4.77 | −0.0463 ± 0.0371 | 0.303 ± 0.486 |
| 01001 | P | I | H | Y | R | 0.141 ± 0.104 | 38.5 ± 3.47 | −0.0871 ± 0.0124 | −0.337 ± 0.621 |
| 00110 | P | T | G | L | H | 0.917 ± 0.0837 | 62.9 ± 4.03 | −0.0321 ± 0.0157 | −0.563 ± 0.945 |
| 00101 | P | T | G | Y | R | 0.283 ± 0.0495 | 62.6 ± 7.20 | −0.0942 ± 0.0122 | −0.415 ± 0.0420 |
| 00011 | P | T | H | L | R | 0.255 ± 0.0835 | 35.8 ± 1.16 | −0.116 ± 0.00717 | −1.26 ± 0.550 |
| 11100 | V | I | G | Y | H | 1.23 ± 0.115 | 57.8 ± 10.5 | −0.0353 ± 0.0229 | −0.0810 ± 0.261 |
| 11010 | V | I | H | L | H | 0.495 ±0.0708 | 52.5 ± 2.52 | −0.0220 ± 0.0362 | 0.0150 ± 0.412 |
| 11001 | V | I | H | Y | R | 0.723 ± 0.146 | 94.4 ± 7.18 | −0.0704 ± 0.0381 | −0.155 ± 0.272 |
| 10110 | V | T | G | L | H | 2.07 ± 0.135 | 71.2 ± 2.26 | −0.0327 ± 0.0395 | −0.0739 ± 0.282 |
| 10101 | V | T | G | Y | R | 0.386 ± 0.0608 | 9.50 ± 0.816 | −0.0696 ± 0.00984 | −0.0435 ± 0.0122 |
| 10011 | V | T | H | L | R | 0.428 ± 0.0405 | 53.4 ± 4.52 | −0.0639 ± 0.0412 | −0.265 ± 0.230 |
| 01110 | P | I | G | L | H | 0.920 ± 0.0284 | 71.1 ± 10.9 | −0.0356 ± 0.0334 | 0.195 ± 0.531 |
| 01101 | P | I | G | Y | R | 0.0835 ± 0.0958 | 38.8 ± 3.98 | −0.0772 ± 0.0279 | −0.112 ± 0.293 |
| 01011 | P | I | H | L | R | 0.126 ± 0.0598 | 22.1 ± 0.712 | −0.120 ± 0.0152 | −0.524 ± 0.175 |
| 00111 | P | T | G | L | R | 3.62 ± 0.380 | 149 ± 4.64 | −0.128 ± 0.0406 | −0.472 ± 0.319 |
| 11110 | V | I | G | L | H | 1.17 ± 0.131 | 43.6 ± 7.23 | −0.0353 ± 0.00761 | −0.00169 ± 0.0987 |
| 11101 | V | I | G | Y | R | 0.100 ± 0.0540 | 8.02 ± 2.70 | −0.0660 ± 0.0277 | −0.0331 ± 0.0182 |
| 11011 | V | I | H | L | R | 0.287 ± 0.0474 | 50.5 ± 3.05 | −0.0659 ± 0.0268 | −0.106 ± 0.0755 |
| 11110 | V | I | G | L | H | 16.5 ± 0.534 | 138 ± 13.7 | −0.0343 ± 0.00316 | −0.0156 ± 0.0830 |
| 01111 | P | I | G | L | R | 3.21 ± 0.176 | 142 ± 22.9 | −0.0808 ± 0.0400 | −0.214 ± 0.637 |
| 11111 | V | I | G | L | R | 26.7 ± 2.68 | 151 ± 6.27 | −0.0427 ± 0.0330 | −0.0559 ± 0.0387 |
| AraC (−) | – | – | – | – | – | 0.128 ± 0.0910 | 0.331 ± 0.0633 | −0.0439 ± 0.0157 | −0.0348 ± 0.0367 |

The difference in fluorescence intensity per OD was calculated from the data provided in Extended Data Fig. 7. Each value corresponds to the mean ± s.d. of three biological replicates.

**Supplementary Table 4 | *Escherichia coli* strains used in this study.**

| Name | Genotype | Source |
|---|---|---|
| JW0063 | F⁻, *Δ(araC–araB)567*, *ΔaraC771::kan*, *ΔlacZ4787*(::rrnB-3), λ⁻, *rph-1*, *Δ(rhaD–rhaB)568*, *hsdR514* | Ref. 55 |
| JW0063 (*Δ*araC) | F⁻, *Δ(araC–araB)567*, *ΔaraC*, *ΔlacZ4787*(::rrnB-3), λ⁻, *rph-1*, *Δ(rhaD–rhaB)568*, *hsdR514* | This study |
| XL10-Gold | Tet$^r$, *ΔmcrA183*, *Δ(mcrCB–hsdSMR–mrr)173*, *endA1*, *supE44*, *thi-1*, *recA1*, *gyrA96*, *relA1*, *lac*, Hte [F', *proAB*, *lacI*$^q$*ZΔM15*, Tn10 (Tet$^r$), Tn5 (Kan$^r$), Amy] | Commercial strain (Agilent Technology, Inc) |

**Supplementary Table 5 | Plasmids used in this study.**

| Name | Based on vector | Origin | Marker | Construct | Source |
|---|---|---|---|---|---|
| pAraC | pET23d | pMB1/ROP | $Amp^R$ | $P_{J23116}$-araC | This study |
| pAraC (−) | | | | (−) | Laboratory stock |
| pBAD7-GFP-TKAPH | pACYC | p15A | $Cm^R$ | $P_{BAD7}$-sfgfp-HSVtk-aph | This study |
| pBAD-GFP-TKAPH | | | | $P_{BAD}$-sfgfp-HSVtk-aph | This study |
| pBAD7/cIO-GFP | | | | $P_{BAD7/OR2}$-sfgfp-HSVtk-aph | This study |
| pAC (−) | | | | (−) | Laboratory stock |
| pBAD7-RFP | pCDF | pCloDF13 | $Strep^R$ | $P_{BAD7}$-mCherry | This study |
| pCDF (−) | | | | (−) | Laboratory stock |
| pBAD-cI | pCDF | pCloDF13* | $Strep^R$ | $P_{BAD}$-lambda cI | This study |

Sequences are provided in Supplementary Note 1.

### Supplementary Table 6 | Primers used in this study.

| Name | Sequence (5' to 3') |
|---|---|
| insert Fwd | GATAAAGCGGGCCATGTTAAGG |
| insert Rev | TCAGCAAAAAACCCCTCAAGACCCGTTTA |
| vector Fwd | TTTTGGATCCCATCATCACCATCAC |
| vector Rev | TTTTCCATGGTAAACCTCCGAGATATAACTAGAGT |
| F15X Fwd | TTTTGGTCTCTCTCGNNKAACGCCCATCTGGTGGCG |
| M42X-I46X Rev | TTTTGGTCTCATGAGMNNATAACCTTTMNNTCCCAGCGGTCGGTCGATAAAAAAATC |
| I46 vector Fwd | TTTTGGTCTCTCTCAATCTCACCATTCGCGGTC |
| F15 vector Rev | TTTTGGTCTCACGAGTATCCCGGCAGCAGG |
| P8P vector Rev | TTGGTCTCTATGGGCGTTAAACGAGTATCCCGGCAGCAGGGGATCATTTTGTGCTTCAGCCATGG |
| P8V vector Rev | TTGGTCTCTATGGGCGTTAAACGAGTATCCCGGCAGCAGCACATCATTTTGTGCTTCAGCCATGG |
| T24T insert Fwd | TTGGTCTCTCCATCTGGTGGCGGGTTTAACGCCGATTGAGGCCAACGGTTATC |
| T24I insert Fwd | TTGGTCTCTCCATCTGGTGGCGGGTTTAATCCCGATTGAGGCCAACGGTTATC |
| H80H-Y82Y insert Rev | TTGGTCTCTGAGCCTCCGGATGACGACCGTAGTGATGAATCTCTCCTGGCGGGAACAG |
| H80G-Y82Y insert Rev | TTGGTCTCTGAGCCTCCGGATGACGACCGTAGTGGCCAATCTCTCCTGGCGGGAACAG |
| H80H-Y82L insert Rev | TTGGTCTCTGAGCCTCCGGATGACGACCCAAGTGATGAATCTCTCCTGGCGGGAACAG |
| H80G-Y82L insert Rev | TTGGTCTCTGAGCCTCCGGATGACGACCCAAGTGGCCAATCTCTCCTGGCGGGAACAG |
| H93H vector Rev | TTGGTCTCTGCTCGCGAATGGTATCACCAGTGGGTTTACTTTCGTCC |
| H93R vector Rev | TTGGTCTCTGCTCGCGAATGGTATCGCCAGTGGGTTTACTTTCGTCC |

**Supplementary Note 1 | Plasmid information.**

| | |
|---|---|
| Name | pAraC |
| Purpose | Directed evolution of AraC. |
| Origin | pMB1/ROP |
| Marker | $Amp^R$ |
| Note | AraC gene is derived from *E. coli str.* K-12 *substr.* MG1655 and fused with additional 24 nt (BamHI-recognition site and histidine tag). |
| Map | 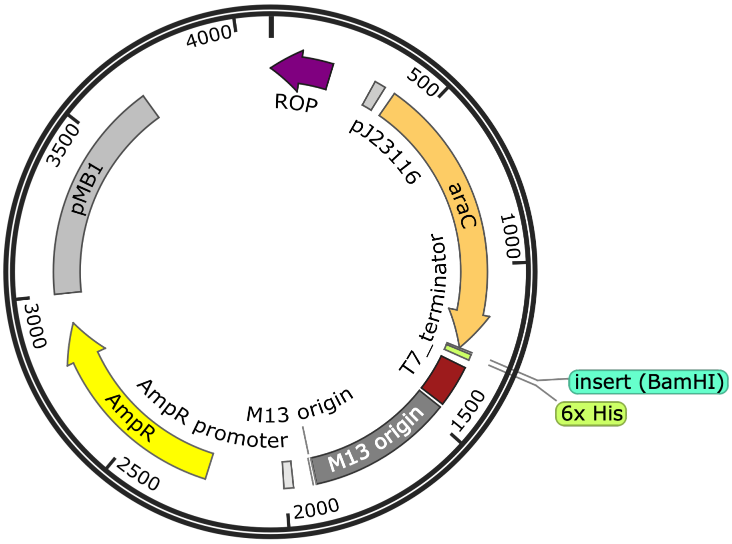 |
| Sequence | TCAGAGGTTTTCACCGTCATCACCGAAACGCGCGAGGCAGCTGCGGTAAAGCTCATCAGCGTGGTCGT<br>GAAGCGATTCACAGATGTCTGCCTGTTCATCCGCGTCCAGCTCGTTGAGTTTCTCCAGAAGCGTTAAT<br>GTCTGGCTTCTGATAAAGCGGGCCATGTTAAGGGCGGTTTTTTCCTGTTTGGTCACTGATGCCTCCGT<br>GTAAGGGGGATTTCTGTTCATGGGGGTAATGATACCGATGAAACGAGAGAGGATGCTCACGATACGGG<br>TTACTGATGATGAACATGCCCGGTTACTGGAACGTTGTGAGGGTAAACAACTGGCGTTGACAGCTAGC<br>TCAGTCCTAGGGACTATGCTAGCTAACAAACTCTAGTTATATCTCGGAGGTTTACCATGGCTGAAGCA<br>CAAAATGATCCCCTGCTGCCGGGATACTCGTTTAACGCCCATCTGGTGGCGGGTTTAACGCCGATTGA<br>GGCCAACGGTTATCTCGATTTTTTTATCGACCGACCGCTGGGAATGAAAGGTTATATTCTCAATCTCA<br>CCATTCGCGGTCAGGGGGTGGTGAAAAATCAGGGACGAGAATTTGTCTGCCGACCGGGTGATATTTTG<br>CTGTTCCCGCCAGGAGAGATTCATCACTACGGTCGTCATCCGGAGGCTCGCGAATGGTATCACCAGTG<br>GGTTTACTTTCGTCCGCGCGCCTACTGGCATGAATGGCTTAACTGGCCGTCAATATTTGCCAATACGG<br>GTTTCTTTCGCCCGGATGAAGCGCACCAGCCGCATTTCAGCGACCTGTTTGGGCAAATCATTAACGCC<br>GGGCAAGGGGAAGGGCGCTATTCGGAGCTGCTGGCGATAAATCTGCTTGAGCAATTGTTACTGCGGCG<br>CATGGAAGCGATTAACGAGTCGCTCCATCCACCGATGGATAATCGGGTACGCGAGGCTTGTCAGTACA<br>TCAGCGATCACCTGGCAGACAGCAATTTTGATATCGCCAGCGTCGCACAGCATGTTTGCTTGTCGCCG<br>TCGCGTCTGTCACATCTTTTCCGCCAGCAGTTAGGGATTAGCGTCTTAAGCTGGCGCGAGGACCAACG<br>CATTAGTCAGGCGAAGCTGCTTTTGAGCACTACCCGGATGCCTATCGCCACCGTCGGTCGCAATGTTG<br>GTTTTGACGATCAACTCTATTTCTCGCGAGTATTTAAAAAATGCACCGGGGCCAGCCCGAGCGAGTTT<br>CGTGCCGGTTGTGAAGAAAAAGTGAATGATGTAGCCGTCAAGTTGTCAGGATCCCATCATCACCATCA<br>CCACTAATGAAAGCTTTAATGAGATCCGGCTGCTAACAAAGCCCGAAAGGAAGCTGAGTTGGCTGCTG<br>CCACCGCTGAGCAATAACTAGCATAACCCCTTGGGGCCTCTAAACGGGTCTTGAGGGGTTTTTTGCTG<br>AAAGGAGGAACTATATCCGGATTGGCGAATGGGACGCGCCCTGTAGCGGCGCATTAAGCGCGGCGGGT<br>GTGGTGGTTACGCGCAGCGTGACCGCTACACTTGCCAGCGCCCTAGCGCCCGCTCCTTTCGCTTTCTT<br>CCCTTCCTTTCTCGCCACGTTCGCCGGCTTTCCCCGTCAAGCTCTAAATCGGGGGCTCCCTTTAGGGT<br>TCCGATTTAGTGCTTTACGGCACCTCGACCCCAAAAAACTTGATTAGGGTGATGGTTCACGTAGTGGG<br>CCATCGCCCTGATAGACGGTTTTTCGCCCTTTGACGTTGGAGTCCACGTTCTTTAATAGTGGACTCTT |

| | |
|---|---|
| | GTTCCAAACTGGAACAACACTCAACCCTATCTCGGTCTATTCTTTTGATTTATAAGGGATTTTGCCGA<br>TTTCGGCCTATTGGTTAAAAAATGAGCTGATTTAACAAAAATTTAACGCGAATTTTAACAAAATATTA<br>ACGCTTACAATTTAGGTGGCACTTTTCGGGGAAATGTGCGCGGAACCCCTATTTGTTTATTTTTCTAA<br>ATACATTCAAATATGTATCCGCTCATGAGACAATAACCCTGATAAATGCTTCAATAATATTGAAAAAG<br>GAAGAGTATGAGTATTCAACATTTCCGTGTCGCCCTTATTCCCTTTTTTGCGGCATTTTGCCTTCCTG<br>TTTTTGCTCACCCAGAAACGCTGGTGAAAGTAAAAGATGCTGAAGATCAGTTGGGTGCACGAGTGGGT<br>TACATCGAACTGGATCTCAACAGCGGTAAGATCCTTGAGAGTTTTCGCCCCGAAGAACGTTTTCCAAT<br>GATGAGCACTTTTAAAGTTCTGCTATGTGGCGCGGTATTATCCCGTATTGACGCCGGGCAAGAGCAAC<br>TCGGTCGCCGCATACACTATTCTCAGAATGACTTGGTTGAGTACTCACCAGTCACAGAAAAGCATCTT<br>ACGGATGGCATGACAGTAAGAGAATTATGCAGTGCTGCCATAACCATGAGTGATAACACTGCGGCCAA<br>CTTACTTCTGACAACGATCGGAGGACCGAAGGAGCTAACCGCTTTTTTGCACAACATGGGGGATCATG<br>TAACTCGCCTTGATCGTTGGGAACCGGAGCTGAATGAAGCCATACCAAACGACGAGCGTGACACCACG<br>ATGCCTGCAGCAATGGCAACAACGTTGCGCAAACTATTAACTGGCGAACTACTTACTCTAGCTTCCCG<br>GCAACAATTAATAGACTGGATGGAGGCGGATAAAGTTGCAGGACCACTTCTGCGCTCGGCCCTTCCGG<br>CTGGCTGGTTTATTGCTGATAAATCTGGAGCCGGTGAGCGTGGTTCTCGCGGTATCATTGCAGCACTG<br>GGGCCAGATGGTAAGCCCTCCCGTATCGTAGTTATCTACACGACGGGGAGTCAGGCAACTATGGATGA<br>ACGAAATAGACAGATCGCTGAGATAGGTGCCTCACTGATTAAGCATTGGTAACTGTCAGACCAAGTTT<br>ACTCATATATACTTTAGATTGATTTAAAACTTCATTTTTAATTTAAAAGGATCTAGGTGAAGATCCTT<br>TTTGATAATCTCATGACCAAAATCCCTTAACGTGAGTTTTCGTTCCACTGAGCGTCAGACCCCGTAGA<br>AAAGATCAAAGGATCTTCTTGAGATCCTTTTTTTCTGCGCGTAATCTGCTGCTTGCAAACAAAAAAAC<br>CACCGCTACCAGCGGTGGTTTGTTTGCCGGATCAAGAGCTACCAACTCTTTTTCCGAAGGTAACTGGC<br>TTCAGCAGAGCGCAGATACCAAATACTGTCCTTCTAGTGTAGCCGTAGTTAGGCCACCACTTCAAGAA<br>CTCTGTAGCACCGCCTACATACCTCGCTCTGCTAATCCTGTTACCAGTGGCTGCTGCCAGTGGCGATA<br>AGTCGTGTCTTACCGGGTTGGACTCAAGACGATAGTTACCGGATAAGGCGCAGCGGTCGGGTTGAACG<br>GGGGGTTCGTGCACACAGCCCAGCTTGGAGCGAACGACCTACACCGAACTGAGATACCTACAGCGTGA<br>GCTATGAGAAAGCGCCACGCTTCCCGAAGGGAGAAAGGCGGACAGGTATCCGGTAAGCGGCAGGGTCG<br>GAACAGGAGAGCGCACGAGGGAGCTTCCAGGGGGAAACGCCTGGTATCTTTATAGTCCTGTCGGGTTT<br>CGCCACCTCTGACTTGAGCGTCGATTTTTGTGATGCTCGTCAGGGGGGCGGAGCCTATGGAAAAACGC<br>CAGCAACGCGGCCTTTTTACGGTTCCTGGCCTTTTGCTGGCCTTTTGCTCACATGTTCTTTCCTGCGT<br>TATCCCCTGATTCTGTGGATAACCGTATTACCGCCTTTGAGTGAGCTGATACCGCTCGCCGCAGCCGA<br>ACGACCGAGCGCAGCGAGTCAGTGAGCGAGGAAGCGGATGAGCGCCTGATGCGGTATTTTCTCCTTAC<br>GCATCTGTGCGGTATTTCACACCGCAATGGTGCACTCTCAGTACAATCTGCTCTGATGCCGCATAGTT<br>AAGCCAGTATACACTCCGCTATCGCTACGTGACTGGGTCATGGCTGCGCCCCGACACCCGCCAACACC<br>CGCTGACGCGCCCTGACGGGCTTGTCTGCTCCCGGCATCCGCTTACAGACAAGCTGTGACCGTCTCCG<br>GGAGCTGCATGTG |

| | |
|---|---|
| Name | pBAD7-GFP-TKAPH |
| Purpose | Reporter and selector plasmid with $P_{BAD7}$ promoter. |
| Origin | p15A |
| Marker | $Cm^R$ |
| Note | $P_{BAD7}$ promoter is designed based on literature[55]. The genes of sfGFP and hsvTK-APH are derived from Waldo et al.[56] and Tominaga et al.[57], respectively. |
| Map | 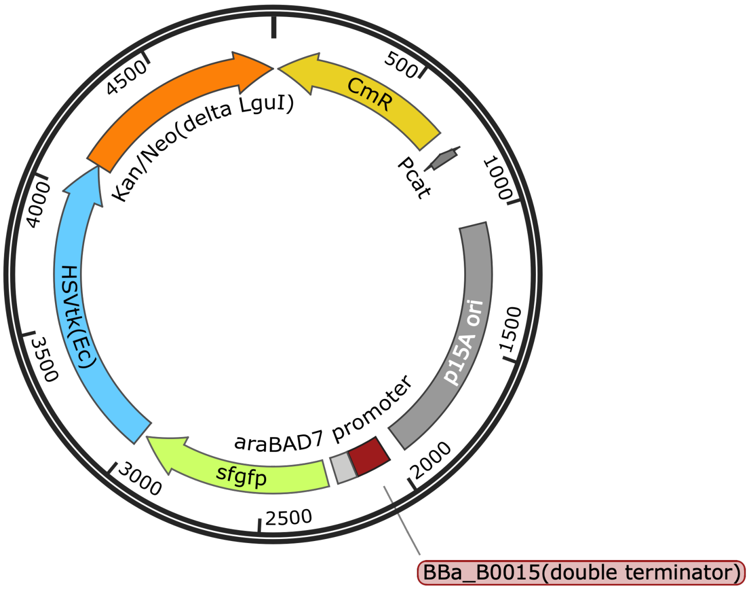 |
| Sequence | CAATAACTGCCTTAAAAAAATTACGCCCCGCCCTGCCACTCATCGCAGTACTGTTGTAATTCATTAAG<br>CATTCTGCCGACATGGAAGCCATCACAAACGGCATGATGAACCTGAATCGCCAGCGGCATCAGCACCT<br>TGTCGCCTTGCGTATAATATTTGCCCATCGTGAAAACGGGGGCGAAGAAGTTGTCCATATTGGCCACG<br>TTTAAATCAAAACTGGTGAAACTCACCCAGGGATTGGCTGAGACGAAAAACATATTCTCAATAAACCC<br>TTTAGGGAAATAGGCCAGGTTTTCACCGTAACACGCCACATCTTGCGAATATATGTGTAGAAACTGCC<br>GGAAATCGTCGTGGTATTCACTCCAGAGCGATGAAAACGTTTCAGTTTGCTCATGGAAAACGGTGTAA<br>CAAGGGTGAACACTATCCCATATCACCAGCTCACCGTCTTTCATTGCCATACGGAATTCCGGATGAGC<br>ATTCATCAGGCGGGCAAGAATGTGAATAAAGGCCGGATAAAACTTGTGCTTATTTTTCTTTACGGTCT<br>TTAAAAAGGCCGTAATATCCAGCTGAACGGTCTGGTTATAGGTACATTGAGCAACTGACTGAAATGCC<br>TCAAAATGTTCTTTACGATGCCATTGGGATATATCAACGGTGGTATATCCAGTGATTTTTTTCTCCAT<br>TTTAGCTTCCTTAGCTCCTGAAAATCTCGATAACTCAAAAAATACGCCCGGTAGTGATCTTATTTCAT<br>TATGGTGAAAGTTGGAACCTCTTACGTGCCGATCAACGTCTCATTTTCGCCAAAAGTTGGCCCAGGGC<br>TTCCCGGTATCAACAGGGACACCAGGATTTATTTATTCTGCGAAGTGATCTTCCGTCACAGGTATTTA<br>TTCGGCGCAAAGTGCGTCGGGTGATGCTGCCAACTTACTGATTTAGTGTATGATGGTGTTTTTGAGGT<br>GCTCCAGTGGCTTCTGTTTCTATCAGCTGTCCCTCCTGTTCAGCTACTGACGGGGTGGTGCGTAACGG<br>CAAAAGCACCGCCGGACATCAGCGCTAGCGGAGTGTATACTGGCTTACTATGTTGGCACTGATGAGGG<br>TGTCAGTGAAGTGCTTCATGTGGCAGGAGAAAAAAGGCTGCACCGGTGCGTCAGCAGAATATGTGATA<br>CAGGATATATTCCGCTTCCTCGCTCACTGACTCGCTACGCTCGGTCGTTCGACTGCGGCGAGCGGAAA<br>TGGCTTACGAACGGGGCGGAGATTTCCTGGAAGATGCCAGGAAGATACTTAACAGGGAAGTGAGAGGG<br>CCGCGGCAAAGCCGTTTTTCCATAGGCTCCGCCCCCCTGACAAGCATCACGAAATCTGACGCTCAAAT<br>CAGTGGTGGCGAAACCCGACAGGACTATAAAGATACCAGGCGTTTCCCCCTGGCGGCTCCCTCGTGCG<br>CTCTCCTGTTCCTGCCTTTCGGTTTACCGGTGTCATTCCGCTGTTATGGCCGCGTTTGTCTCATTCCA<br>CGCCTGACACTCAGTTCCGGGTAGGCAGTTCGCTCCAAGCTGGACTGTATGCACGAACCCCCCGTTCA<br>GTCCGACCGCTGCGCCTTATCCGGTAACTATCGTCTTGAGTCCAACCCGGAAAGACATGCAAAAGCAC<br>CACTGGCAGCAGCCACTGGTAATTGATTTAGAGGAGTTAGTCTTGAAGTCATGCGCCGGTTAAGGCTA<br>AACTGAAAGGACAAGTTTTGGTGACTGCGCTCCTCCAAGCCAGTTACCTCGGTTCAAAGAGTTGGTAG<br>CTCAGAGAACCTTCGAAAAACCGCCCTGCAAGGCGGTTTTTTCGTTTTCAGAGCAAGAGATTACGCGC<br>AGACCAAAACGATCTCAAGAAGATCATCTTATTAATCAGATAAAATATTTCAAGATTTCAGTGCAATT<br>TATCTCTTCAAATGTAGCACCTGAAGTCAGCCCCATACGATATAAGTTGTAATTCTCATGTTTGACAG<br>CTTATCATCGATCACTGATAGTGCTAGTGTAGATCACTACTAGAGCCAGGCATCAAATAAAACGAAAG |

| | |
|---|---|
| | GCTCAGTCGAAAGACTGGGCCTTTCGTTTTATCTGTTGTTTGTCGGTGAACGCTCTCTACTAGAGTCA<br>CACTGGCTCACCTTCGGGTGGGCCTTTCTGCGTTTATATACTAGCATTTTTATCCATAAGATTAGCAT<br>TTTTATCCATAGATCCTGGTACCGAATTCATTGTCTACTGTTTCTATAACACGGGATTAAAAATACAA<br>GGAGAAATTCAATGGGGTCTAAAGGCGAAGAACTGTTCACCGGCGTAGTTCCGATCCTGGTTGAACTG<br>GACGGTGACGTTAATGGTCATAAGTTCTCTGTTCGTGGTGAAGGTGAGGGCGACGCGACCAACGGTAA<br>ACTGACCCTGAAATTCATCTGCACCACTGGCAAACTGCCGGTTCCGTGGCCGACTCTGGTTACCACCC<br>TGACCTATGGTGTTCAGTGCTTCTCTCGTTACCCGGATCACATGAAACAGCACGACTTCTTCAAATCT<br>GCGATGCCGGAGGGTTATGTTCAGGAACGTACCATCTCTTTCAAGGATGACGGCACCTACAAAACCCG<br>TGCCGAAGTTAAATTCGAGGGTGATACGCTGGTAAACCGCATCGAACTGAAAGGTATCGACTTCAAAG<br>AGGACGGTAATATCCTCGGTCACAAGCTGGAATACAACTTCAACTCTCACAACGTTTACATCACCGCG<br>GACAAACAGAAAAACGGTATCAAAGCGAACTTTAAGATCCGTCACAATGTTGAAGACGGCAGCGTTCA<br>GCTCGCTGACCACTACCAACAAAATACCCCGATTGGCGACGGTCCGGTTCTGCTGCCGGACAACCACT<br>ATCTGTCTACCCAGTCTGTGCTCTCTAAGGACCCGAACGAGAAACGTGACCACATGGTGCTGCTGGAG<br>TTCGTGACCGCAGCGGGCATCACGCACGGCATGGACGAACTGTACAAATGATTAAATCTCAGCAGGTC<br>CCCATGGCGAGCTATCCGGGTCACCAGCATGCATCTGCTTTCGATCAGGCAGCGCGCAGCCGTGGTCA<br>TTCTAATCGTCGTACCGCACTGCGTCCGCGTCGTCAGCAGGAGGCCACTGAGGTTCGTCCGGAGCAAA<br>AGATGCCGACCCTGTTACGCGTATACATTGATGGCCGCATGGTATGGGTAAAACCACCACGACCCAA<br>TTACTGGTTGCGCTGGGCAGCCGTGATGATATTGTTTATGTGCCTGAACCGATGACGTATTGGCGCGT<br>GCTGGGCGCGAGTGAAACTATTGCTAATATCTATACGACCCAGCATCGTCTGGACCAAGGGGAAATCA<br>GCGCCGGTGATGCAGCCGTAGTGATGACCAGTGCGCAAATCACGATGGGTATGCCTTACGCAGTAACC<br>GATGCGGTTCTGGCGCCGCATATTGGTGGTGAAGCCGGCAGTAGCCATGCGCCGCCGCCTGCCCTGAC<br>CCTGATTTTTGATCGTCACCCGATTGCGGCTCTGCTGTGCTATCCTGCTGCACGTTATCTGATGGGTT<br>CTATGACCCCACAGGCCGTCCTGGCATTCGTTGCACTGATTCCGCCTACTCTGCCTGGGACCAATATC<br>GTGCTGGGGGCGCTGCCAGAAGATCGTCATATCGACCGTCTGGCGAAACGTCAACGTCCTGGTGAACG<br>CCTGGATCTGGCGATGCTGGCAGCGATTCGTCGTGTATATGGCCTGCTGGCGAACACTGTCCGTTACC<br>TGCAATGCGGTGGCAGTTGGCGTGAAGATTGGGGTCAACTGAGCGGTACGGCAGTTCCTCCGCAGGGT<br>GCGGAACCTCAGTCTAACGCAGGTCCGCGTCCGCACATTGGTGATACCCTGTTCACCCTGTTCCGTGC<br>GCCGGAGCTGCTGGCACCAAATGGGGATCTGTACAATGTTTTCGCGTGGGCGCTGGATGTTCTGGCTA<br>AGCGTCTGCGCAGCATGCATGTTTTTATTCTGGATTATGATCAAAGCCCAGCAGGCTGTCGTGATGCG<br>CTGCTTCAACTGACTAGCGGCATGGTGCAAACGCATGTGACGACGCCTGGGAGTATCCCGACCATCTG<br>TGATCTTGCCCGTACCTTCGCACGTGAAATGGGTGAAGCGAATATTGAACAAGATGGATTGCACGCAG<br>GTTCTCCGGCCGCTTGGGTGGAGAGGCTATTCGGCTATGACTGGGCACAACAGACAATCGGCTGCTCT<br>GATGCCGCCGTGTTCCGGCTGTCAGCGCAGGGGCGCCCGGTTCTTTTTGTCAAGACCGACCTGTCCGG<br>TGCCCTGAATGAACTGCAGGACGAGGCAGCGCGGCTATCGTGGCTGGCCACGACGGGCGTTCCTTGCG<br>CAGCTGTGCTCGACGTTGTCACTGAAGCGGGAAGGGACTGGCTGCTATTGGGCGAAGTGCCGGGGCAG<br>GATCTCCTGTCATCTCACCTTGCTCCTGCCGAGAAAGTATCCATCATGGCTGATGCAATGCGGCGGCT<br>GCATACGCTTGATCCGGCTACCTGCCCATTCGACCACCAAGCGAAACATCGCATCGAGCGAGCACGTA<br>CTCGGATGGAAGCCGGTCTTGTCGATCAGGATGATCTGGACGAAGTGCATCAGGGGCTCGCGCCAGCC<br>GAACTGTTCGCCAGGCTCAAGGCGCGCATGCCCGACGGCGAGGATCTCGTCGTGACCCATGGCGATGC<br>CTGCTTGCCGAATATCATGGTGGAAAATGGCCGCTTTTCTGGATTCATCGACTGTGGCCGGCTGGGTG<br>TGGCGGACCGCTATCAGGACATAGCGTTGGCTACCCGTGATATTGCTGAAGTGCTTGGCGGCGAATGG<br>GCTGACCGCTTCCTCGTGCTTTACGGTATCGCCGCTCCCGATTCGCAGCGCATCGCCTTCTATCGCCT<br>TCTTGACGAGTTCTTCTGA |

| | |
|---|---|
| Name | pBAD-GFP-TKAPH |
| Purpose | Reporter and selector plasmid with $P_{BAD}$ promoter. |
| Origin | p15A |
| Marker | $Cm^R$ |
| Note | $P_{BAD}$ promoter sequence is obtained from BioBrick I746908. |
| Map | 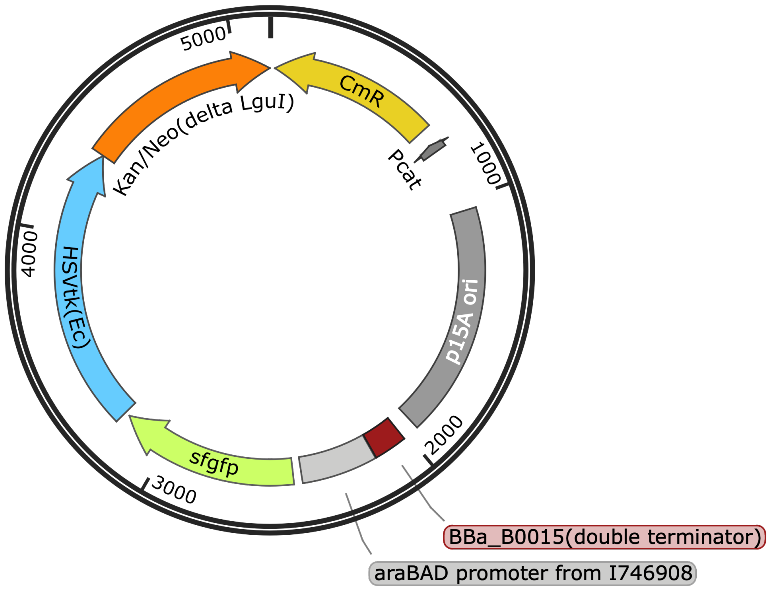 |
| Sequence | CAATAACTGCCTTAAAAAAATTACGCCCCGCCCTGCCACTCATCGCAGTACTGTTGTAATTCATTAAG<br>CATTCTGCCGACATGGAAGCCATCACAAACGGCATGATGAACCTGAATCGCCAGCGGCATCAGCACCT<br>TGTCGCCTTGCGTATAATATTTGCCCATCGTGAAAACGGGGGCGAAGAAGTTGTCCATATTGGCCACG<br>TTTAAATCAAAACTGGTGAAACTCACCCAGGGATTGGCTGAGACGAAAAACATATTCTCAATAAACCC<br>TTTAGGGAAATAGGCCAGGTTTTCACCGTAACACGCCACATCTTGCGAATATATGTGTAGAAACTGCC<br>GGAAATCGTCGTGGTATTCACTCCAGAGCGATGAAAACGTTTCAGTTTGCTCATGGAAAACGGTGTAA<br>CAAGGGTGAACACTATCCCATATCACCAGCTCACCGTCTTTCATTGCCATACGGAATTCCGGATGAGC<br>ATTCATCAGGCGGGCAAGAATGTGAATAAAGGCCGGATAAAACTTGTGCTTATTTTTCTTTACGGTCT<br>TTAAAAAGGCCGTAATATCCAGCTGAACGGTCTGGTTATAGGTACATTGAGCAACTGACTGAAATGCC<br>TCAAAATGTTCTTTACGATGCCATTGGGATATATCAACGGTGGTATATCCAGTGATTTTTTTCTCCAT<br>TTTAGCTTCCTTAGCTCCTGAAAATCTCGATAACTCAAAAAATACGCCCGGTAGTGATCTTATTTCAT<br>TATGGTGAAAGTTGGAACCTCTTACGTGCCGATCAACGTCTCATTTTCGCCAAAAGTTGGCCCAGGGC<br>TTCCCGGTATCAACAGGGACACCAGGATTTATTTATTCTGCGAAGTGATCTTCCGTCACAGGTATTTA<br>TTCGGCGCAAAGTGCGTCGGGTGATGCTGCCAACTTACTGATTTAGTGTATGATGGTGTTTTTGAGGT<br>GCTCCAGTGGCTTCTGTTTCTATCAGCTGTCCCTCCTGTTCAGCTACTGACGGGGTGGTGCGTAACGG<br>CAAAAGCACCGCCGGACATCAGCGCTAGCGGAGTGTATACTGGCTTACTATGTTGGCACTGATGAGGG<br>TGTCAGTGAAGTGCTTCATGTGGCAGGAGAAAAAAGGCTGCACCGGTGCGTCAGCAGAATATGTGATA<br>CAGGATATATTCCGCTTCCTCGCTCACTGACTCGCTACGCTCGGTCGTTCGACTGCGGCGAGCGGAAA<br>TGGCTTACGAACGGGGCGGAGATTTCCTGGAAGATGCCAGGAAGATACTTAACAGGGAAGTGAGAGGG<br>CCGCGGCAAAGCCGTTTTTCCATAGGCTCCGCCCCCCTGACAAGCATCACGAAATCTGACGCTCAAAT<br>CAGTGGTGGCGAAACCCGACAGGACTATAAAGATACCAGGCGTTTCCCCCTGGCGGCTCCCTCGTGCG<br>CTCTCCTGTTCCTGCCTTTCGGTTTACCGGTGTCATTCCGCTGTTATGGCCGCGTTTGTCTCATTCCA<br>CGCCTGACACTCAGTTCCGGGTAGGCAGTTCGCTCCAAGCTGGACTGTATGCACGAACCCCCCGTTCA<br>GTCCGACCGCTGCGCCTTATCCGGTAACTATCGTCTTGAGTCCAACCCGGAAAGACATGCAAAAGCAC<br>CACTGGCAGCAGCCACTGGTAATTGATTTAGAGGAGTTAGTCTTGAAGTCATGCGCCGGTTAAGGCTA<br>AACTGAAAGGACAAGTTTTGGTGACTGCGCTCCTCCAAGCCAGTTACCTCGGTTCAAAGAGTTGGTAG<br>CTCAGAGAACCTTCGAAAAACCGCCCTGCAAGGCGGTTTTTTCGTTTTCAGAGCAAGAGATTACGCGC<br>AGACCAAAACGATCTCAAGAAGATCATCTTATTAATCAGATAAAATATTTCAAGATTTCAGTGCAATT<br>TATCTCTTCAAATGTAGCACCTGAAGTCAGCCCCATACGATATAAGTTGTAATTCTCATGTTTGACAG<br>CTTATCATCGATCACTGATAGTGCTAGTGTAGATCACTACTAGAGCCAGGCATCAAATAAAACGAAAG<br>GCTCAGTCGAAAGACTGGGCCTTTCGTTTTATCTGTTGTTTGTCGGTGAACGCTCTCTACTAGAGTCA |

| | |
|---|---|
| | CACTGGCTCACCTTCGGGTGGGCCTTTCTGCGTTTATATACATTCAGAGAAGAAACCAATTGTCCATA<br>TTGCATCAGACATTGCCGTCACTGCGTCTTTTACTGGCTCTTCTCGCTAACCAAACCGGTAACCCCGC<br>TTATTAAAAGCATTCTGTAACAAAGCGGGACCAAAGCCATGACAAAAACGCGTAACAAAAGTGTCTAT<br>AATCACGGCAGAAAAGTCCACATTGATTATTTGCACGGCGTCACACTTTGCTATGCCATAGCATTTTT<br>ATCCATAAGATTAGCGGATCCTACCTGACGCTTTTTATCGCAACTCTCTACTGTTTCTCCATATAACA<br>CGGGATTAAAAATACAAGGAGAAATTCAATGGGGTCTAAAGGCGAAGAACTGTTCACCGGCGTAGTTC<br>CGATCCTGGTTGAACTGGACGGTGACGTTAATGGTCATAAGTTCTCTGTTCGTGGTGAAGGTGAGGGC<br>GACGCGACCAACGGTAAACTGACCCTGAAATTCATCTGCACCACTGGCAAACTGCCGGTTCCGTGGCC<br>GACTCTGGTTACCACCCTGACCTATGGTGTTCAGTGCTTCTCTCGTTACCCGGATCACATGAAACAGC<br>ACGACTTCTTCAAATCTGCGATGCCGGAGGGTTATGTTCAGGAACGTACCATCTCTTTCAAGGATGAC<br>GGCACCTACAAAACCCGTGCCGAAGTTAAATTCGAGGGTGATACGCTGGTAAACCGCATCGAACTGAA<br>AGGTATCGACTTCAAAGAGGACGGTAATATCCTCGGTCACAAGCTGGAATACAACTTCAACTCTCACA<br>ACGTTTACATCACCGCGGACAAACAGAAAAACGGTATCAAAGCGAACTTTAAGATCCGTCACAATGTT<br>GAAGACGGCAGCGTTCAGCTCGCTGACCACTACCAACAAAATACCCCGATTGGCGACGGTCCGGTTCT<br>GCTGCCGGACAACCACTATCTGTCTACCCAGTCTGTGCTCTCTAAGGACCCGAACGAGAAACGTGACC<br>ACATGGTGCTGCTGGAGTTCGTGACCGCAGCGGGCATCACGCACGGCATGGACGAACTGTACAAATGA<br>TTAAATCTCAGCAGGTCCCCATGGCGAGCTATCCGGGTCACCAGCATGCATCTGCTTTCGATCAGGCA<br>GCGCGCAGCCGTGGTCATTCTAATCGTCGTACCGCACTGCGTCCGCGTCGTCAGCAGGAGGCCACTGA<br>GGTTCGTCCGGAGCAAAAGATGCCGACCCTGTTACGCGTATACATTGATGGGCCGCATGGTATGGGTA<br>AAACCACCACGACCCAATTACTGGTTGCGCTGGGCAGCCGTGATGATATTGTTTATGTGCCTGAACCG<br>ATGACGTATTGGCGCGTGCTGGGCGCGAGTGAAACTATTGCTAATATCTATACGACCCAGCATCGTCT<br>GGACCAAGGGGAAATCAGCGCCGGTGATGCAGCCGTAGTGATGACCAGTGCGCAAATCACGATGGGTA<br>TGCCTTACGCAGTAACCGATGCGGTTCTGGCGCCGCATATTGGTGGTGAAGCCGGCAGTAGCCATGCG<br>CCGCCGCCTGCCCTGACCCTGATTTTTGATCGTCACCCGATTGCGGCTCTGCTGTGCTATCCTGCTGC<br>ACGTTATCTGATGGGTTCTATGACCCCACAGGCCGTCCTGGCATTCGTTGCACTGATTCCGCCTACTC<br>TGCCTGGGACCAATATCGTGCTGGGGGCGCTGCCAGAAGATCGTCATATCGACCGTCTGGCGAAACGT<br>CAACGTCCTGGTGAACGCCTGGATCTGGCGATGCTGGCAGCGATTCGTCGTGTATATGGCCTGCTGGC<br>GAACACTGTCCGTTACCTGCAATGCGGTGGCAGTTGGCGTGAAGATTGGGGTCAACTGAGCGGTACGG<br>CAGTTCCTCCGCAGGGTGCGGAACCTCAGTCTAACGCAGGTCCGCGTCCGCACATTGGTGATACCCTG<br>TTCACCCTGTTCCGTGCGCCGGAGCTGCTGGCACCAAATGGGGATCTGTACAATGTTTTCGCGTGGGC<br>GCTGGATGTTCTGGCTAAGCGTCTGCGCAGCATGCATGTTTTTATTCTGGATTATGATCAAAGCCCAG<br>CAGGCTGTCGTGATGCGCTGCTTCAACTGACTAGCGGCATGGTGCAAACGCATGTGACGACGCCTGGG<br>AGTATCCCGACCATCTGTGATCTTGCCCGTACCTTCGCACGTGAAATGGGTGAAGCGAATATTGAACA<br>AGATGGATTGCACGCAGGTTCTCCGGCCGCTTGGGTGGAGAGGCTATTCGGCTATGACTGGGCACAAC<br>AGACAATCGGCTGCTCTGATGCCGCCGTGTTCCGGCTGTCAGCGCAGGGGCGCCCGGTTCTTTTTGTC<br>AAGACCGACCTGTCCGGTGCCCTGAATGAACTGCAGGACGAGGCAGCGCGGCTATCGTGGCTGGCCAC<br>GACGGGCGTTCCTTGCGCAGCTGTGCTCGACGTTGTCACTGAAGCGGGAAGGGACTGGCTGCTATTGG<br>GCGAAGTGCCGGGGCAGGATCTCCTGTCATCTCACCTTGCTCCTGCCGAGAAAGTATCCATCATGGCT<br>GATGCAATGCGGCGGCTGCATACGCTTGATCCGGCTACCTGCCCATTCGACCACCAAGCGAAACATCG<br>CATCGAGCGAGCACGTACTCGGATGGAAGCCGGTCTTGTCGATCAGGATGATCTGGACGAAGTGCATC<br>AGGGGCTCGCGCCAGCCGAACTGTTCGCCAGGCTCAAGGCGCGCATGCCCGACGGCGAGGATCTCGTC<br>GTGACCCATGGCGATGCCTGCTTGCCGAATATCATGGTGGAAAATGGCCGCTTTTCTGGATTCATCGA<br>CTGTGGCCGGCTGGGTGTGGCGGACCGCTATCAGGACATAGCGTTGGCTACCCGTGATATTGCTGAAG<br>TGCTTGGCGGCGAATGGGCTGACCGCTTCCTCGTGCTTTACGGTATCGCCGCTCCCGATTCGCAGCGC<br>ATCGCCTTCTATCGCCTTCTTGACGAGTTCTTCTGA |

| | |
|---|---|
| Name | pBAD7-RFP |
| Purpose | Reporter plasmid that activates mCherry expression under $P_{BAD7}$ promoter. |
| Origin | pCloDF13 |
| Marker | $Strep^R$ |
| Note | Gene of mCherry is obtained from BioBrick J06504. |
| Map | 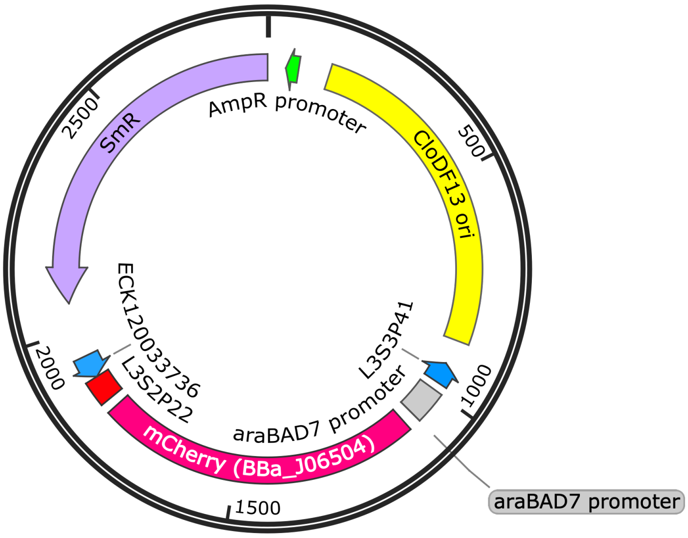 |
| Sequence | ACTCTTCCTTTTTCAATATTATTGAAGCATTTATCAGGGTTATTGTCTCATGAGCGGATACATATTTG<br>AATGTATTTAGAAAAATAAACAAATAGCTAGCTCACTCGGTCGCTACGCTCCGGGCGTGAGACTGCGG<br>CGGGCGCTGCGGACACATACAAAGTTACCCACAGATTCCGTGGATAAGCAGGGGACTAACATGTGAGG<br>CAAAACAGCAGGGCCGCGCCGGTGGCGTTTTTCCATAGGCTCCGCCCTCCTGCCAGAGTTCACATAAA<br>CAGACGCTTTTCCGGTGCATCTGTGGGAGCCGTGAGGCTCAACCATGAATCTGACAGTACGGGCGAAA<br>CCCGACAGGACTTAAAGATCCCCACCGTTTCCGGCGGGTCGCTCCCTCTTGCGCTCTCCTGTTCCGAC<br>CCTGCCGTTTACCGGATACCTGTTCCGCCTTTCTCCCTTACGGGAAGTGTGGCGCTTTCTCATAGCTC<br>ACACACTGGTATCTCGGCTCGGTGTAGGTCGTTCGCTCCAAGCTGGGCTGTAAGCAAGAACTCCCCGT<br>TCAGCCCGACTGCTGCGCCTTATCCGGTAACTGTTCACTTGAGTCCAACCCGGAAAAGCACGGTAAAA<br>CGCCACTGGCAGCAGCCATTGGTAACTGGGAGTTCGCAGAGGATTTGTTTAGCTAAACACGCGGTTGC<br>TCTTGAAGTGTGCGCCAAAGTCCGGCTACACTGGAAGGACAGATTTGGTTGCTGTGCTCTGCGAAAGC<br>CAGTTACCACGGTTAAGCAGTTCCCCAACTGACTTAACCTTCGATCAAACCACCTCCCCAGGTGGTTT<br>TTTCGTTTACAGGGCAAAAGATTACGCGCAGAAAAAAAGGATCTCAAGAAGATCCTTTGATCTTTTCT<br>ACTGAACCGCTCTAGATTTCAGTGCAATTTATCTCTTCAAATGAGGATCCAAAAAAAAAAAACACCCT<br>AACGGGTGTTTTTTTTTTTTTGGTCTGCCTGACACCATGCAAGCTTTAGCATTTTTATCCATAAGATT<br>AGCATTTTTATCCATAGATCCTGGTACCGAATTCATTGTCTACTGTTTCTATAAATTTGAGAGGGGAG<br>CAACTAGTATGGTTAGCAAGGGCGAGGAGGATAACATGGCCATCATCAAGGAGTTCATGCGCTTCAAG<br>GTGCACATGGAGGGCTCCGTGAACGGCCACGAGTTCGAGATCGAGGGCGAGGGCGAGGGCCGCCCCTA<br>CGAGGGCACCCAGACCGCCAAGCTGAAGGTGACCAAGGGTGGCCCCCTGCCCTTCGCCTGGGACATCC<br>TGTCCCCTCAGTTCATGTACGGCTCCAAGGCCTACGTGAAGCACCCCGCCGACATCCCCGACTACTTG<br>AAGCTGTCCTTCCCCGAGGGCTTCAAGTGGGAGCGCGTGATGAACTTCGAGGACGGCGGCGTGGTGAC<br>CGTGACCCAGGACTCCTCCTTGCAGGACGGCGAGTTCATCTACAAGGTGAAGCTGCGCGGCACCAACT<br>TCCCCTCCGACGGCCCCGTAATGCAGAAGAAGACCATGGGCTGGGAGGCCTCCTCCGAGCGGATGTAC<br>CCCGAGGACGGCGCCCTGAAGGGCGAGATCAAGCAGAGGCTGAAGCTGAAGGACGGCGGCCACTACGA<br>CGCTGAGGTCAAGACCACCTACAAGGCCAAGAAGCCCGTGCAGCTGCCCGGCGCCTACAACGTCAACA<br>TCAAGTTGGACATCACCTCCCACAACGAGGACTACACCATCGTGGAACAGTACGAACGCGCCGAGGGC<br>CGCCACTCCACCGGCGGCATGGACGAGCTGTACAAGTAGTAACATATGTCAGTCGACGAGCTCGGTAC<br>CAAATTCCAGAAAAGAGGCCGCGAAAGCGGCCTTTTTTCGTTTTGGTCCGCGCAATAAAAAAGCCCCC<br>GGAAGGTGATCTTCCGGGGGCTTTCTCATGCGTTACAATATCGTTCACTCATCCAATCTCACTGACGG<br>TTAGGCGTGCCTGGTACCCTGCCCTGAACCGACGACCGGGTCATCGTGGCCGGATCTTGCGGCCCCTC<br>GGCTTGAACGAATTGTTAGACATTATTTGCCGACTACCTTGGTGATCTCGCCTTTCACGTAGTGGACA<br>AATTCTTCCAACTGATCTGCGCGCGAGGCCAAGCGATCTTCTTCTTGTCCAAGATAAGCCTGTCTAGC<br>TTCAAGTATGACGGGCTGATACTGGGCCGGCAGGCGCTCCATTGCCCAGTCGGCAGCGACATCCTTCG |

| | |
|---|---|
| | GCGCGATTTTGCCGGTTACTGCGCTGTACCAAATGCGGGACAACGTAAGCACTACATTTCGCTCATCG<br>CCAGCCCAGTCGGGCGGCGAGTTCCATAGCGTTAAGGTTTCATTTAGCGCCTCAAATAGATCCTGTTC<br>AGGAACCGGATCAAAGAGTTCCTCCGCCGCTGGACCTACCAAGGCAACGCTATGTTCTCTTGCTTTTG<br>TCAGCAAGATAGCCAGATCAATGTCGATCGTGGCTGGCTCGAAGATACCTGCAAGAATGTCATTGCGC<br>TGCCATTCTCCAAATTGCAGTTCGCGCTTAGCTGGATAACGCCACGGAATGATGTCGTCGTGCACAAC<br>AATGGTGACTTCTACAGCGCGGAGAATCTCGCTCTCTCCAGGGGAAGCCGAAGTTTCCAAAAGGTCGT<br>TGATCAAAGCTCGCCGCGTTGTTTCATCAAGCCTTACGGTCACCGTAACCAGCAAATCAATATCACTG<br>TGTGGCTTCAGGCCGCCATCCACTGCGGAGCCGTACAAATGTACGGCCAGCAACGTCGGTTCGAGATG<br>GCGCTCGATGACGCCAACTACCTCTGATAGTTGAGTCGATACTTCGGCGATCACCGCTTCCCTCAT |

<table>
<tr><td>Name</td><td>pBAD7/cIO-GFP</td></tr>
<tr><td>Purpose</td><td>Integration of output from $P_{BAD}$ promoter toward $P_{BAD7}$ promoter via lambda cI repressor.</td></tr>
<tr><td>Origin</td><td>p15A</td></tr>
<tr><td>Marker</td><td>$Cm^R$</td></tr>
<tr><td>Note</td><td>One of the lambda cI binding sites, OR2, was inserted to $P_{BAD7}$ promoter.</td></tr>
<tr><td>Map</td><td>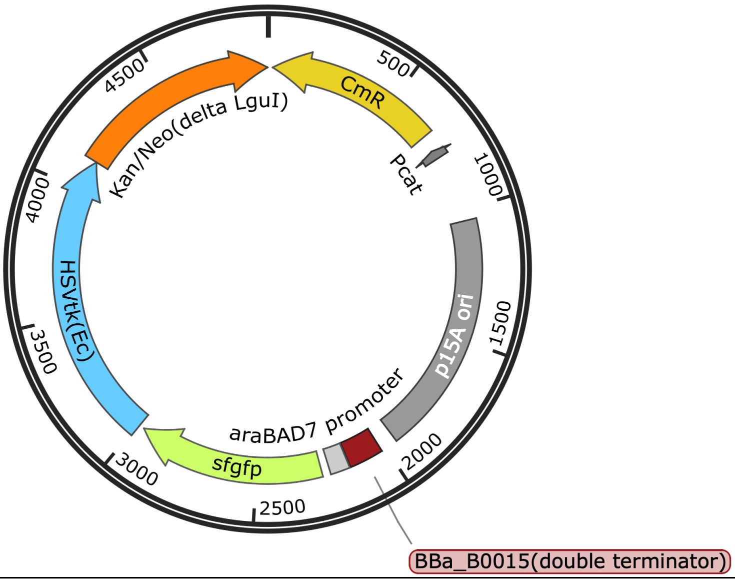
</td></tr>
<tr><td>Sequence</td><td>CAATAACTGCCTTAAAAAAATTACGCCCCGCCCTGCCACTCATCGCAGTACTGTTGTAATTCATTAAG<br>CATTCTGCCGACATGGAAGCCATCACAAACGGCATGATGAACCTGAATCGCCAGCGGCATCAGCACCT<br>TGTCGCCTTGCGTATAATATTTGCCCATCGTGAAAACGGGGGCGAAGAAGTTGTCCATATTGGCCACG<br>TTTAAATCAAAACTGGTGAAACTCACCCAGGGATTGGCTGAGACGAAAAACATATTCTCAATAAACCC<br>TTTAGGGAAATAGGCCAGGTTTTCACCGTAACACGCCACATCTTGCGAATATATGTGTAGAAACTGCC<br>GGAAATCGTCGTGGTATTCACTCCAGAGCGATGAAAACGTTTCAGTTTGCTCATGGAAAACGGTGTAA<br>CAAGGGTGAACACTATCCCATATCACCAGCTCACCGTCTTTCATTGCCATACGGAATTCCGGATGAGC<br>ATTCATCAGGCGGGCAAGAATGTGAATAAAGGCCGGATAAAACTTGTGCTTATTTTTCTTTACGGTCT<br>TTAAAAAGGCCGTAATATCCAGCTGAACGGTCTGGTTATAGGTACATTGAGCAACTGACTGAAATGCC<br>TCAAAATGTTCTTTACGATGCCATTGGGATATATCAACGGTGGTATATCCAGTGATTTTTTTCTCCAT<br>TTTAGCTTCCTTAGCTCCTGAAAATCTCGATAACTCAAAAAATACGCCCGGTAGTGATCTTATTTCAT<br>TATGGTGAAAGTTGGAACCTCTTACGTGCCGATCAACGTCTCATTTTCGCCAAAAGTTGGCCCAGGGC<br>TTCCCGGTATCAACAGGGACACCAGGATTTATTTATTCTGCGAAGTGATCTTCCGTCACAGGTATTTA<br>TTCGGCGCAAAGTGCGTCGGGTGATGCTGCCAACTTACTGATTTAGTGTATGATGGTGTTTTTGAGGT<br>GCTCCAGTGGCTTCTGTTTCTATCAGCTGTCCCTCCTGTTCAGCTACTGACGGGGTGGTGCGTAACGG<br>CAAAAGCACCGCCGGACATCAGCGCTAGCGGAGTGTATACTGGCTTACTATGTTGGCACTGATGAGGG<br>TGTCAGTGAAGTGCTTCATGTGGCAGGAGAAAAAAGGCTGCACCGGTGCGTCAGCAGAATATGTGATA<br>CAGGATATATTCCGCTTCCTCGCTCACTGACTCGCTACGCTCGGTCGTTCGACTGCGGCGAGCGGAAA<br>TGGCTTACGAACGGGGCGGAGATTTCCTGGAAGATGCCAGGAAGATACTTAACAGGGAAGTGAGAGGG<br>CCGCGGCAAAGCCGTTTTTCCATAGGCTCCGCCCCCCTGACAAGCATCACGAAATCTGACGCTCAAAT<br>CAGTGGTGGCGAAACCCGACAGGACTATAAAGATACCAGGCGTTTCCCCCTGGCGGCTCCCTCGTGCG<br>CTCTCCTGTTCCTGCCTTTCGGTTTACCGGTGTCATTCCGCTGTTATGGCCGCGTTTGTCTCATTCCA<br>CGCCTGACACTCAGTTCCGGGTAGGCAGTTCGCTCCAAGCTGGACTGTATGCACGAACCCCCCGTTCA<br>GTCCGACCGCTGCGCCTTATCCGGTAACTATCGTCTTGAGTCCAACCCGGAAAGACATGCAAAAGCAC<br>CACTGGCAGCAGCCACTGGTAATTGATTTAGAGGAGTTAGTCTTGAAGTCATGCGCCGGTTAAGGCTA<br>AACTGAAAGGACAAGTTTTGGTGACTGCGCTCCTCCAAGCCAGTTACCTCGGTTCAAAGAGTTGGTAG<br>CTCAGAGAACCTTCGAAAAACCGCCCTGCAAGGCGGTTTTTTCGTTTTCAGAGCAAGAGATTACGCGC<br>AGACCAAAACGATCTCAAGAAGATCATCTTATTAATCAGATAAAATATTTCAAGATTTCAGTGCAATT<br>TATCTCTTCAAATGTAGCACCTGAAGTCAGCCCCATACGATATAAGTTGTAATTCTCATGTTTGACAG<br>CTTATCATCGATCACTGATAGTGCTAGTGTAGATCACTACTAGAGCCAGGCATCAAATAAAACGAAAG<br>GCTCAGTCGAAAGACTGGGCCTTTCGTTTTATCTGTTGTTTGTCGGTGAACGCTCTCTACTAGAGTCA</td></tr>
</table>

| | |
|---|---|
| | CACTGGCTCACCTTCGGGTGGGCCTTTCTGCGTTTATATACTAGCATTTTTATCCATAAGATTAGCAT<br>TTTTATCCATAGATCCTAACACCGTGCGTGTTGTCTACTGTTTCTATAACCACGGTTTTAATAAGGAG<br>GTATTCATATGGGGTCTAAAGGCGAAGAACTGTTCACCGGCGTAGTTCCGATCCTGGTTGAACTGGAC<br>GGTGACGTTAATGGTCATAAGTTCTCTGTTCGTGGTGAAGGTGAGGGCGACGCGACCAACGGTAAACT<br>GACCCTGAAATTCATCTGCACCACTGGCAAACTGCCGGTTCCGTGGCCGACTCTGGTTACCACCCTGA<br>CCTATGGTGTTCAGTGCTTCTCTCGTTACCCGGATCACATGAAACAGCACGACTTCTTCAAATCTGCG<br>ATGCCGGAGGGTTATGTTCAGGAACGTACCATCTCTTTCAAGGATGACGGCACCTACAAAACCCGTGC<br>CGAAGTTAAATTCGAGGGTGATACGCTGGTAAACCGCATCGAACTGAAAGGTATCGACTTCAAAGAGG<br>ACGGTAATATCCTCGGTCACAAGCTGGAATACAACTTCAACTCTCACAACGTTTACATCACCGCGGAC<br>AAACAGAAAAACGGTATCAAAGCGAACTTTAAGATCCGTCACAATGTTGAAGACGGCAGCGTTCAGCT<br>CGCTGACCACTACCAACAAAATACCCCGATTGGCGACGGTCCGGTTCTGCTGCCGGACAACCACTATC<br>TGTCTACCCAGTCTGTGCTCTCTAAGGACCCGAACGAGAAACGTGACCACATGGTGCTGCTGGAGTTC<br>GTGACCGCAGCGGGCATCACGCACGGCATGGACGAACTGTACAAATGATTAAATCTCAGCAGGTCCCC<br>ATGGCGAGCTATCCGGGTCACCAGCATGCATCTGCTTTCGATCAGGCAGCGCGCAGCCGTGGTCATTC<br>TAATCGTCGTACCGCACTGCGTCCGCGTCGTCAGCAGGAGGCCACTGAGGTTCGTCCGGAGCAAAAGA<br>TGCCGACCCTGTTACGCGTATACATTGATGGGCCGCATGGTATGGGTAAAACCACCACGACCCAATTA<br>CTGGTTGCGCTGGGCAGCCGTGATGATATTGTTTATGTGCCTGAACCGATGACGTATTGGCGCGTGCT<br>GGGCGCGAGTGAAACTATTGCTAATATCTATACGACCCAGCATCGTCTGGACCAAGGGGAAATCAGCG<br>CCGGTGATGCAGCCGTAGTGATGACCAGTGCGCAAATCACGATGGGTATGCCTTACGCAGTAACCGAT<br>GCGGTTCTGGCGCCGCATATTGGTGGTGAAGCCGGCAGTAGCCATGCGCCGCCGCCTGCCCTGACCCT<br>GATTTTTGATCGTCACCCGATTGCGGCTCTGCTGTGCTATCCTGCTGCACGTTATCTGATGGGTTCTA<br>TGACCCCACAGGCCGTCCTGGCATTCGTTGCACTGATTCCGCCTACTCTGCCTGGGACCAATATCGTG<br>CTGGGGGCGCTGCCAGAAGATCGTCATATCGACCGTCTGGCGAAACGTCAACGTCCTGGTGAACGCCT<br>GGATCTGGCGATGCTGGCAGCGATTCGTCGTGTATATGGCCTGCTGGCGAACACTGTCCGTTACCTGC<br>AATGCGGTGGCAGTTGGCGTGAAGATTGGGGTCAACTGAGCGGTACGGCAGTTCCTCCGCAGGGTGCG<br>GAACCTCAGTCTAACGCAGGTCCGCGTCCGCACATTGGTGATACCCTGTTCACCCTGTTCCGTGCGCC<br>GGAGCTGCTGGCACCAAATGGGGATCTGTACAATGTTTTCGCGTGGGCGCTGGATGTTCTGGCTAAGC<br>GTCTGCGCAGCATGCATGTTTTTATTCTGGATTATGATCAAAGCCCAGCAGGCTGTCGTGATGCGCTG<br>CTTCAACTGACTAGCGGCATGGTGCAAACGCATGTGACGACGCCTGGGAGTATCCCGACCATCTGTGA<br>TCTTGCCCGTACCTTCGCACGTGAAATGGGTGAAGCGAATATTGAACAAGATGGATTGCACGCAGGTT<br>CTCCGGCCGCTTGGGTGGAGAGGCTATTCGGCTATGACTGGGCACAACAGACAATCGGCTGCTCTGAT<br>GCCGCCGTGTTCCGGCTGTCAGCGCAGGGGCGCCCGGTTCTTTTTGTCAAGACCGACCTGTCCGGTGC<br>CCTGAATGAACTGCAGGACGAGGCAGCGCGGCTATCGTGGCTGGCCACGACGGGCGTTCCTTGCGCAG<br>CTGTGCTCGACGTTGTCACTGAAGCGGGAAGGGACTGGCTGCTATTGGGCGAAGTGCCGGGGCAGGAT<br>CTCCTGTCATCTCACCTTGCTCCTGCCGAGAAAGTATCCATCATGGCTGATGCAATGCGGCGGCTGCA<br>TACGCTTGATCCGGCTACCTGCCCATTCGACCACCAAGCGAAACATCGCATCGAGCGAGCACGTACTC<br>GGATGGAAGCCGGTCTTGTCGATCAGGATGATCTGGACGAAGTGCATCAGGGGCTCGCGCCAGCCGAA<br>CTGTTCGCCAGGCTCAAGGCGCGCATGCCCGACGGCGAGGATCTCGTCGTGACCCATGGCGATGCCTG<br>CTTGCCGAATATCATGGTGGAAAATGGCCGCTTTTCTGGATTCATCGACTGTGGCCGGCTGGGTGTGG<br>CGGACCGCTATCAGGACATAGCGTTGGCTACCCGTGATATTGCTGAAGTGCTTGGCGGCGAATGGGCT<br>GACCGCTTCCTCGTGCTTTACGGTATCGCCGCTCCCGATTCGCAGCGCATCGCCTTCTATCGCCTTCT<br>TGACGAGTTCTTCTGA |

| | |
|---|---|
| Name | pBAD-cI |
| Purpose | Inverting the output of $P_{BAD}$ promoter. |
| Origin | pCloDF13 |
| Marker | $Strep^R$ |
| Note | G56A mutation in the CloDF13 origin, highlighted in yellow in the “Sequence” section, seems to increase the copy number (about 30 copies/cell). |
| Map | 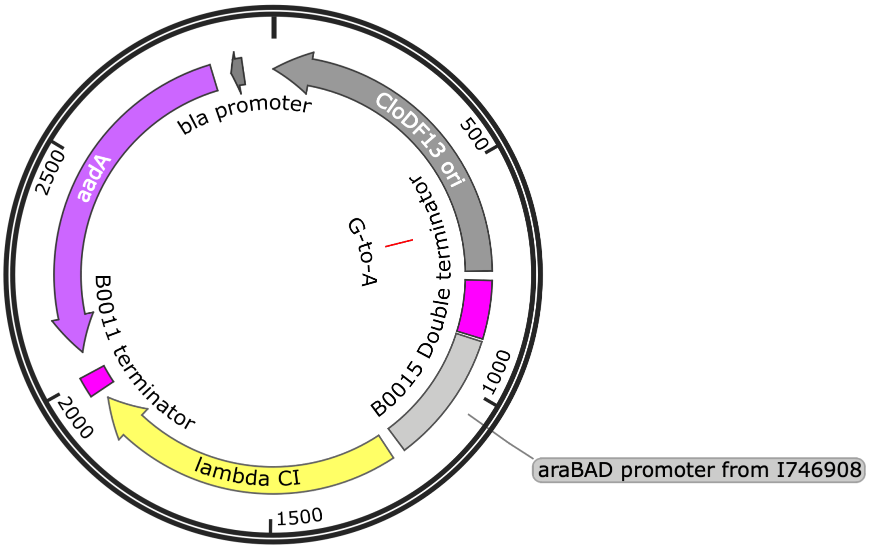 |
| Sequence | GCGCTGCGGACACATACAAAGTTACCCACAGATTCCGTGGATAAGCAGGGGACTAACATGTGAGGCAA<br>AACAGCAGGGCCGCGCCGGTGGCGTTTTTCCATAGGCTCCGCCCTCCTGCCAGAGTTCACATAAACAG<br>ACGCTTTTCCGGTGCATCTGTGGGAGCCGTGAGGCTCAACCATGAATCTGACAGTACGGGCGAAACCC<br>GACAGGACTTAAAGATCCCCACCGTTTCCGGCGGGTCGCTCCCTCTTGCGCTCTCCTGTTCCGACCCT<br>GCCGTTTACCGGATACCTGTTCCGCCTTTCTCCCTTACGGGAAGTGTGGCGCTTTCTCATAGCTCACA<br>CACTGGTATCTCGGCTCGGTGTAGGTCGTTCGCTCCAAGCTGGGCTGTAAGCAAGAACTCCCCGTTCA<br>GCCCGACTGCTGCGCCTTATCCGGTAACTGTTCACTTGAGTCCAACCCGGAAAAGCACGGTAAAACGC<br>CACTGGCAGCAGCCATTGGTAACTGGGAGTTCGCAGAGGATTTGTTTAGCTAAACACGCGGTTGCTCT<br>TGAAGTGTGCGCCAAAGTCCGGCTACACTGGAAGGACAGATTTGGTTGCTGTGCTCTGCGAAAGCCAG<br>TTACCACGGTTAAGCAGTTTCCCAACTGACTTAACCTTCGATCAAACCACCTCCCCAGGTGGTTTTTT<br>CGTTTACAGGGCAAAAGATTACGCGCAGAAAAAAAGGATCTCAAGAAGATCCTTTGATCTTTTCTACT<br>GAACCGCTCTAGCCAGGCATCAAATAAAACGAAAGGCTCAGTCGAAAGACTGGGCCTTTCGTTTTATC<br>TGTTGTTTGTCGGTGAACGCTCTCTACTAGAGTCACACTGGCTCACCTTCGGGTGGGCCTTTCTGCGT<br>TTATAGGTACCATTCAGAGAAGAAACCAATTGTCCATATTGCATCAGACATTGCCGTCACTGCGTCTT<br>TTACTGGCTCTTCTCGCTAACCAAACCGGTAACCCCGCTTATTAAAAGCATTCTGTAACAAAGCGGGA<br>CCAAAGCCATGACAAAAACGCGTAACAAAAGTGTCTATAATCACGGCAGAAAAGTCCACATTGATTAT<br>TTGCACGGCGTCACACTTTGCTATGCCATAGCATTTTTATCCATAAGATTAGCGGATCCTACCTGACG<br>CTTTTTATCGCAACTCTCTACTGTTTCTCCATATAACGTGAGTTGTTGGGGGTGTGCATCAGATGAGC<br>ACAAAAAGAAACCATTAACACAAGAGCAGCTTGAGGACGCACGTCGCCTTAAAGCAATTTATGAAAA<br>AAAGAAAAATGAACTTGGCTTATCCCAGGAATCTGTCGCAGACAAGATGGGGATGGGGCAGTCAGGCG<br>TTGGTGCTTTATTTAATGGCATCAATGCATTAAATGCTTATAACGCCGCATTGCTTGCAAAAATTCTC<br>AAAGTTAGCGTTGAAGAATTTAGCCCTTCAATCGCCAGAGAAATCTACGAGATGTATGAAGCGGTTAG<br>TATGCAGCCGTCACTTAGAAGTGAGTATGAGTACCCTGTTTTTTCTCATGTTCAGGCAGGGATGTTCT<br>CACCTGAGCTTAGAACCTTTACCAAAGGTGATGCGGAGAGATGGGTAAGCACAACCAAAAAAGCCAGT<br>GATTCTGCATTCTGGCTTGAGGTTGAAGGTAATTCCATGACCGCACCAACAGGCTCCAAGCCAAGCTT<br>TCCTGACGGAATGTTAATTCTCGTTGACCCTGAGCAGGCTGTTGAGCCAGGTGATTTCTGCATAGCCA<br>GACTTGGGGGTGATGAGTTTACCTTCAAGAAACTGATCAGGGATAGCGGTCAGGTGTTTTTACAACCA<br>CTAAACCCACAGTACCCAATGATCCCATGCAATGAGAGTTGTTCCGTTGTGGGGAAAGTTATCGCTAG<br>TCAGTGGCCTGAAGAGACGTTTGGCGCTGCATAATAAGCTTACTAGTaataCTGCAGAGAGAATATAA<br>AAAGCCAGATTATTAATCCGGCTTTTTATTATTTGTCATCGTGGCCGGATCTTGCGGCCCCTCGGCT<br>TGAACGAATTGTTAGACATTATTTGCCGACTACCTTGGTGATCTCGCCTTTCACGTAGTGGACAAATT |

| | |
|---|---|
| | CTTCCAACTGATCTGCGCGCGAGGCCAAGCGATCTTCTTCTTGTCCAAGATAAGCCTGTCTAGCTTCA<br>AGTATGACGGGCTGATACTGGGCCGGCAGGCGCTCCATTGCCCAGTCGGCAGCGACATCCTTCGGCGC<br>GATTTTGCCGGTTACTGCGCTGTACCAAATGCGGGACAACGTAAGCACTACATTTCGCTCATCGCCAG<br>CCCAGTCGGGCGGCGAGTTCCATAGCGTTAAGGTTTCATTTAGCGCCTCAAATAGATCCTGTTCAGGA<br>ACCGGATCAAAGAGTTCCTCCGCCGCTGGACCTACCAAGGCAACGCTATGTTCTCTTGCTTTTGTCAG<br>CAAGATAGCCAGATCAATGTCGATCGTGGCTGGCTCGAAGATACCcGCAAGAATGTCATTGCGCTGCC<br>ATTCTCCAAATTGCAGTTCGCGCTTAGCTGGATAACGCCAaGGAATGATGTCGTCGTGCACAACAATG<br>GTGACTTCTACAGCGCGGAGAATCTCGCTCTCTCCAGGGGAAGCCGAAGTTTCCAAAAGGTCGTTGAT<br>CAAAGCTCGCCGCGTTGTTTCATCAAGCCTTACGGTCACCGTAACCAGCAAATCAATATCACTGTGTG<br>GCTTCAGGCCGCCATCCACTGCGGAGCCGTACAAATGTACGGCCAGCAACGTCGGTTCGAGATGGCGC<br>TCGATGACGCCAACTACCTCTGATAGTTGAGTCGATACTTCGGCGATCACCGCTTCCCTCATACTCTT<br>CCTTTTTCAATATTATTGAAGCATTTATCAGGGTTATTGTCTCATGAGCGGATACATATTTGAATGTA<br>TTTAGAAAAATAAACAAATAGCTAGCTCACTCGGTCGCTACGCTCCGGGCGTGAGACTGCGGCGG |

| | |
|---|---|
| Name | pAraC (−) |
| Purpose | Negative control for AraC. |
| Origin | pMB1/ROP |
| Marker | $Amp^R$ |
| Note | |
| Map | 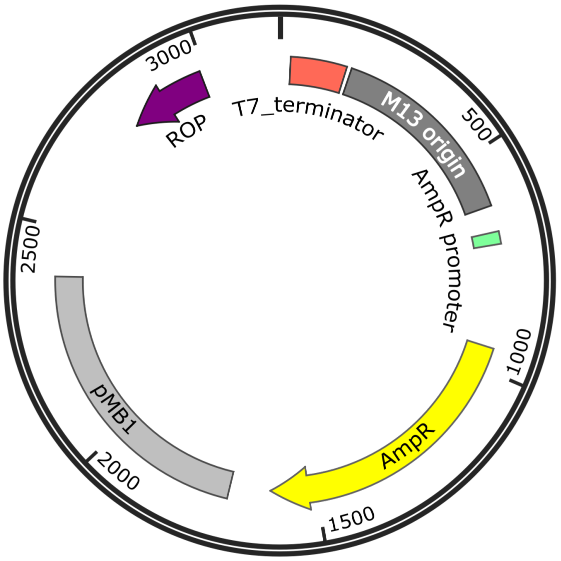 |
| Sequence | TTTTGCTCTTCATAATGAGATCCGGCTGCTAACAAAGCCCGAAAGGAAGCTGAGTTGGCTGCTGCCAC<br>CGCTGAGCAATAACTAGCATAACCCCTTGGGGCCTCTAAACGGGTCTTGAGGGGTTTTTTGCTGAAAG<br>GAGGAACTATATCCGGATTGGCGAATGGGACGCGCCCTGTAGCGGCGCATTAAGCGCGGCGGGTGTGG<br>TGGTTACGCGCAGCGTGACCGCTACACTTGCCAGCGCCCTAGCGCCCGCTCCTTTCGCTTTCTTCCCT<br>TCCTTTCTCGCCACGTTCGCCGGCTTTCCCCGTCAAGCTCTAAATCGGGGGCTCCCTTTAGGGTTCCG<br>ATTTAGTGCTTTACGGCACCTCGACCCCAAAAAACTTGATTAGGGTGATGGTTCACGTAGTGGGCCAT<br>CGCCCTGATAGACGGTTTTTCGCCCTTTGACGTTGGAGTCCACGTTCTTTAATAGTGGACTCTTGTTC<br>CAAACTGGAACAACACTCAACCCTATCTCGGTCTATTCTTTTGATTTATAAGGGATTTTGCCGATTTC<br>GGCCTATTGGTTAAAAAATGAGCTGATTTAACAAAAATTTAACGCGAATTTTAACAAAATATTAACGC<br>TTACAATTTAGGTGGCACTTTTCGGGGAAATGTGCGCGGAACCCCTATTTGTTTATTTTTCTAAATAC<br>ATTCAAATATGTATCCGCTCATGAGACAATAACCCTGATAAATGCTTCAATAATATTGAAAAAGGAAG<br>AGTATGAGTATTCAACATTTCCGTGTCGCCCTTATTCCCTTTTTTGCGGCATTTTGCCTTCCTGTTTT<br>TGCTCACCCAGAAACGCTGGTGAAAGTAAAAGATGCTGAAGATCAGTTGGGTGCACGAGTGGGTTACA<br>TCGAACTGGATCTCAACAGCGGTAAGATCCTTGAGAGTTTTCGCCCCGAAGAACGTTTTCCAATGATG<br>AGCACTTTTAAAGTTCTGCTATGTGGCGCGGTATTATCCCGTATTGACGCCGGGCAAGAGCAACTCGG<br>TCGCCGCATACACTATTCTCAGAATGACTTGGTTGAGTACTCACCAGTCACAGAAAAGCATCTTACGG<br>ATGGCATGACAGTAAGAGAATTATGCAGTGCTGCCATAACCATGAGTGATAACACTGCGGCCAACTTA<br>CTTCTGACAACGATCGGAGGACCGAAGGAGCTAACCGCTTTTTTGCACAACATGGGGGATCATGTAAC<br>TCGCCTTGATCGTTGGGAACCGGAGCTGAATGAAGCCATACCAAACGACGAGCGTGACACCACGATGC<br>CTGCAGCAATGGCAACAACGTTGCGCAAACTATTAACTGGCGAACTACTTACTCTAGCTTCCCGGCAA<br>CAATTAATAGACTGGATGGAGGCGGATAAAGTTGCAGGACCACTTCTGCGCTCGGCCCTTCCGGCTGG<br>CTGGTTTATTGCTGATAAATCTGGAGCCGGTGAGCGTGGGTCTCGCGGTATCATTGCAGCACTGGGGC<br>CAGATGGTAAGCCCTCCCGTATCGTAGTTATCTACACGACGGGGAGTCAGGCAACTATGGATGAACGA<br>AATAGACAGATCGCTGAGATAGGTGCCTCACTGATTAAGCATTGGTAACTGTCAGACCAAGTTTACTC<br>ATATATACTTTAGATTGATTTAAAACTTCATTTTTAATTTAAAAGGATCTAGGTGAAGATCCTTTTTG<br>ATAATCTCATGACCAAAATCCCTTAACGTGAGTTTTCGTTCCACTGAGCGTCAGACCCCGTAGAAAAG<br>ATCAAAGGATCTTCTTGAGATCCTTTTTTTCTGCGCGTAATCTGCTGCTTGCAAACAAAAAAACCACC<br>GCTACCAGCGGTGGTTTGTTTGCCGGATCAAGAGCTACCAACTCTTTTTCCGAAGGTAACTGGCTTCA<br>GCAGAGCGCAGATACCAAATACTGTCCTTCTAGTGTAGCCGTAGTTAGGCCACCACTTCAAGAACTCT<br>GTAGCACCGCCTACATACCTCGCTCTGCTAATCCTGTTACCAGTGGCTGCTGCCAGTGGCGATAAGTC<br>GTGTCTTACCGGGTTGGACTCAAGACGATAGTTACCGGATAAGGCGCAGCGGTCGGGTTGAACGGGGG<br>GTTCGTGCACACAGCCCAGCTTGGAGCGAACGACCTACACCGAACTGAGATACCTACAGCGTGAGCTA<br>TGAGAAAGCGCCACGCTTCCCGAAGGGAGAAAGGCGGACAGGTATCCGGTAAGCGGCAGGGTCGGAAC<br>AGGAGAGCGCACGAGGGAGCTTCCAGGGGGAAACGCCTGGTATCTTTATAGTCCTGTCGGGTTTCGCC<br>ACCTCTGACTTGAGCGTCGATTTTTGTGATGCTCGTCAGGGGGGCGGAGCCTATGGAAAAACGCCAGC |

| | |
|---|---|
| | AACGCGGCCTTTTTACGGTTCCTGGCCTTTTGCTGGCCTTTTGCTCACATGTTCTTTCCTGCGTTATC<br>CCCTGATTCTGTGGATAACCGTATTACCGCCTTTGAGTGAGCTGATACCGCTCGCCGCAGCCGAACGA<br>CCGAGCGCAGCGAGTCAGTGAGCGAGGAAGCGGATGAGCGCCTGATGCGGTATTTTCTCCTTACGCAT<br>CTGTGCGGTATTTCACACCGCAATGGTGCACTCTCAGTACAATCTGCTCTGATGCCGCATAGTTAAGC<br>CAGTATACACTCCGCTATCGCTACGTGACTGGGTCATGGCTGCGCCCCGACACCCGCCAACACCCGCT<br>GACGCGCCCTGACGGGCTTGTCTGCTCCCGGCATCCGCTTACAGACAAGCTGTGACCGTCTCCGGGAG<br>CTGCATGTGTCAGAGGTTTTCACCGTCATCACCGAAACGCGCGAGGCAGCTGCGGTAAAGCTCATCAG<br>CGTGGTCGTGAAGCGATTCACAGATGTCTGCCTGTTCATCCGCGTCCAGCTCGTTGAGTTTCTCCAGA<br>AGCGTTAATGTCTGGCTTCTGATAAAGCGGGCCATGTTAAGGGCGGTTTTTTCCTGTTTGGTCACTGA<br>TGCCTCCGTGTAAGGGGGATTTCTGTTCATGGGGGTAATGATACCGATGAAACGAGAGAGGATGCTCA<br>CGATACGGGTTACTGATGATGAACATGCCCGGTTACTGGAACGTTGTGAGGGTAAACAACTGGCGTAA<br>CACCGTGCGTGTTGACTATTTTACCTCTGGCGGAGAAGAGCAAAA |

DNA sequences are written in the 5' to 3' direction and colors represent target gene promoters in blue, putative ribosome binding sites in orange, protein coding sequences in red, and plasmid backbone in black. All RBS sequences for target genes were designed by using RBS calculator[58]. Plasmid maps were depicted by SnapGene Viewer.
